# Mutant-*SETBP1* activates transcription of Myc programs to accelerate *CSF3R*-driven myeloproliferative neoplasms

**DOI:** 10.1101/2021.11.04.467367

**Authors:** Sarah A. Carratt, Garth L. Kong, Brittany M. Smith, Zachary Schonrock, Lauren Maloney, Adrian Baris, Brian J. Druker, Theodore P. Braun, Julia E. Maxson

**Author notes:** **Correspondence** Julia E. Maxson, Knight Cancer Institute, Oregon Health & Science University, Mail code: KR-HEM, 3181 S.W. Sam Jackson Park Rd., Portland, OR 97239.

## Abstract

Colony stimulating factor 3 receptor (*CSF3R*) mutations lead to JAK pathway activation and are the molecular hallmark of chronic neutrophilic leukemia (CNL). Approximately half of CNL patients also have mutations in SET binding protein 1 (*SETBP1)*. In this study, we developed models of *SETBP1*-mutant leukemia to understand the role that SETBP1 plays in CNL. *SETBP1* mutations promote self-renewal of *CSF3R-*mutant hematopoietic progenitors *in vitro* and prevent cells from undergoing terminal differentiation. *In vivo, SETBP1* mutations accelerate leukemia progression, leading to the rapid development of hepatosplenomegaly and granulocytosis. Through transcriptomic and epigenomic profiling, we found that *SETBP1* enhances progenitor-associated programs—most strongly upregulating *Myc* and Myc target genes. This upregulation of *Myc* can be reversed by epigenetic modulatory drugs. In summary, we find that *SETBP1* mutations promote aggressive hematopoietic cell expansion when expressed with mutant *CSF3R* through the upregulation of Myc-associated gene expression programs.

**Statement of Significance:** *SETBP1* is frequently mutated in chronic neutrophilic leukemia, but its role in the biology of this disease is unclear. We find that mutant *SETBP1* enhances transcription of *Myc* and Myc target genes to promote aggressive disease biology, and that these oncogenic transcriptional programs can be reversed by epigenetic modulatory drugs.

## Introduction

Chronic neutrophilic leukemia (CNL) is a rare myeloproliferative neoplasm characterized by the overproduction of neutrophils. Colony stimulating factor 3 receptor (*CSF3R*) mutations are the molecular hallmark of CNL and lead to ligand-independent receptor dimerization and downstream janus activated kinase (JAK) pathway activation^1^. Historically, treatment options for CNL were limited. The discovery of activating *CSF3R* mutations in CNL led to the identification of JAK inhibitors as a potential targeted therapeutic strategy for these patients. In a clinical trial for patients with CNL and atypical CML a 54% overall response rate was achieved with the JAK1/2 inhibitor, ruxolitinib, in those patients who had mutations in *CSF3R*^2^. Although targeting CSF3R signaling with ruxolitinib has shown clinical efficacy, responses are not always durable. Anecdotally, the small number of long-term responders tended to have less genetic complexity. Treatment of CNL will therefore likely require a multipronged therapeutic approach to improve initial treatment response rates and prevent the development of acquired resistance.

One of the most commonly co-mutated genes in CNL is SET binding protein 1 (*SETBP1)*—which is mutated in approximately half of these leukemias^2^. In myeloid leukemia, *SETBP1* mutations (*SETBP1*^*Mut*^) predominantly occur in the β-TrCP degron motif. One of the two most common point mutations, D868N, is used in these studies. *SETBP1* point mutations interfere with the ubiquitination and subsequent degradation of SETBP1, resulting in an accumulation of SETBP1^Mut^ protein^3^. Mutations in *SETBP1* are often associated with poor prognosis in myeloid malignancies^4^, however high levels of wildtype *SETBP1* (*SETBP1*^*Wt*^) also drives adverse outcomes in acute myeloid leukemia^5^.

SETBP1 regulates tumor suppressor pathways and modulates transcription^6-9^. SETBP1 is a binding partner of SET, a 39-kDa protein that inhibits the tumor suppressor protein phosphatase 2A (PP2A)^10^. Setbp1 has also been implicated as a transcriptional regulator in murine leukemia models, conferring increased self-renewal capacity through enhanced expression of *Hoxa9, Hoxa10*, and *Myb* and repression of *Runx1* expression^7,*9,11*^. In a human embryonic kidney model (Flp-In™-293), SETBP1 was shown to recruit the MLL1 transcriptional activator complex and directly upregulate *MECOM* and MECOM target genes^8^. Recently, we found that SETBP1 can modulate disease biology driven by co-occurring mutations^12^. Specifically, in the context of Ras-pathway driven leukemia, mutant SETBP1 can increase MAPK pathway activation^12^. The goal of this study was to understand the context specific role of *SETBP1* mutations in CNL to enable the development of therapeutic approaches that improve treatment outcomes for these patients.

In this study, we investigate how *SETBP1* modulates *CSF3R*-driven disease biology. In a murine model of *CSF3R*-driven CNL, we find that the addition of a *SETBP1* mutation enhances cellular proliferation and accelerates disease progression. In a cell line expressing mutant *SETBP1*, we find that one of the strongest proliferation-associated signatures is that of MYC target genes. Expression of mutant *SETBP1* both increases *Myc* gene expression and activates a MYC E-box luciferase reporter. When we assessed SETBP1-driven histone modulation, we identified a 67% overlap between Myc binding sites and H3K4me3 marks upregulated by SETBP1, indicating an overlap in the promoters that are regulated by Myc and SETBP1. Treatment with LSD1 inhibitors decreased *Myc* expression by at least 70% for each of the three inhibitors evaluated (GSK2879552, GSK-LSD1, ORY-1001). LSD1 inhibitors caused synergistic cell death when combined with the JAK inhibitor, ruxolitinib. As a mutation that drives robust proliferation in our model systems, *SETBP1* represents a promising candidate for targeted therapeutic development.

## Results

One of the primary goals of this study was to understand how the presence of a *SETBP1* mutation alters *CSF3R*-driven phenotypes in both murine and *in vitro* models. To understand how mutant *SETBP1* modulates the phenotypes associated with a *CSF3R* point mutation (T618I), we first performed a murine hematopoietic colony forming unit (CFU) assay. In this assay, primary mouse bone marrow cells were transduced with retroviral vectors to express our mutations of interest and 5,000 sorted cells per condition were plated in cytokine-free methylcellulose. Interestingly, while *CSF3R*^*T618I*^ expressed alongside an empty vector control leads to the formation of large dispersed colonies, neither *SETBP1*^*Wt*^ or *SETBP1*^*D868N*^ with empty vector stimulate any colony formation in the absence of cytokines (**Figure 1A**). When combined with *CSF3R*^*T618I*^, overexpression of *SETBP1* (either *SETBP1*^*Wt*^ or *SETBP1*^*D868N*^) significantly augmented colony formation and the colonies had large dense centers (**Figure 1A,B)**. This augmentation by *SETBP1*^*Wt*^ driven off a strong promoter is consistent with the known mechanism of *SETBP1*^*D868N*^ driving oncogenesis through protein overexpression. Cytospins prepared from individual colonies showed that they are primarily composed of myeloid cells (**Figure 1C**). To determine whether expressing both oncogenes conferred replating potential, colonies were harvested, washed with PBS, and approximately 10,000 cells were resuspended in fresh cytokine-free methylcellulose. Both *CSF3R*^*T618I*^+*SETBP1*^*Wt*^ and *CSF3R*^*T618I*^+*SETBP1*^*D868N*^ expression in cells confers re-plating potential out to at least the fourth passage in CFU assay (**Figure 1D**). To determine if *SETBP1*^*D868N*^ can augment proliferation driven by activation of endogenous CSF3R, cells expressing either *SETBP1*^*D868N*^ or an empty vector were plated in methylcellulose with 100 nM GSCF, the ligand for CSF3R. In this assay, GCSF-driven colony formation increased by a factor of 6 when *SETBP1*^*D868N*^ was expressed (**Figure 1E,F**).

**Figure 1.**
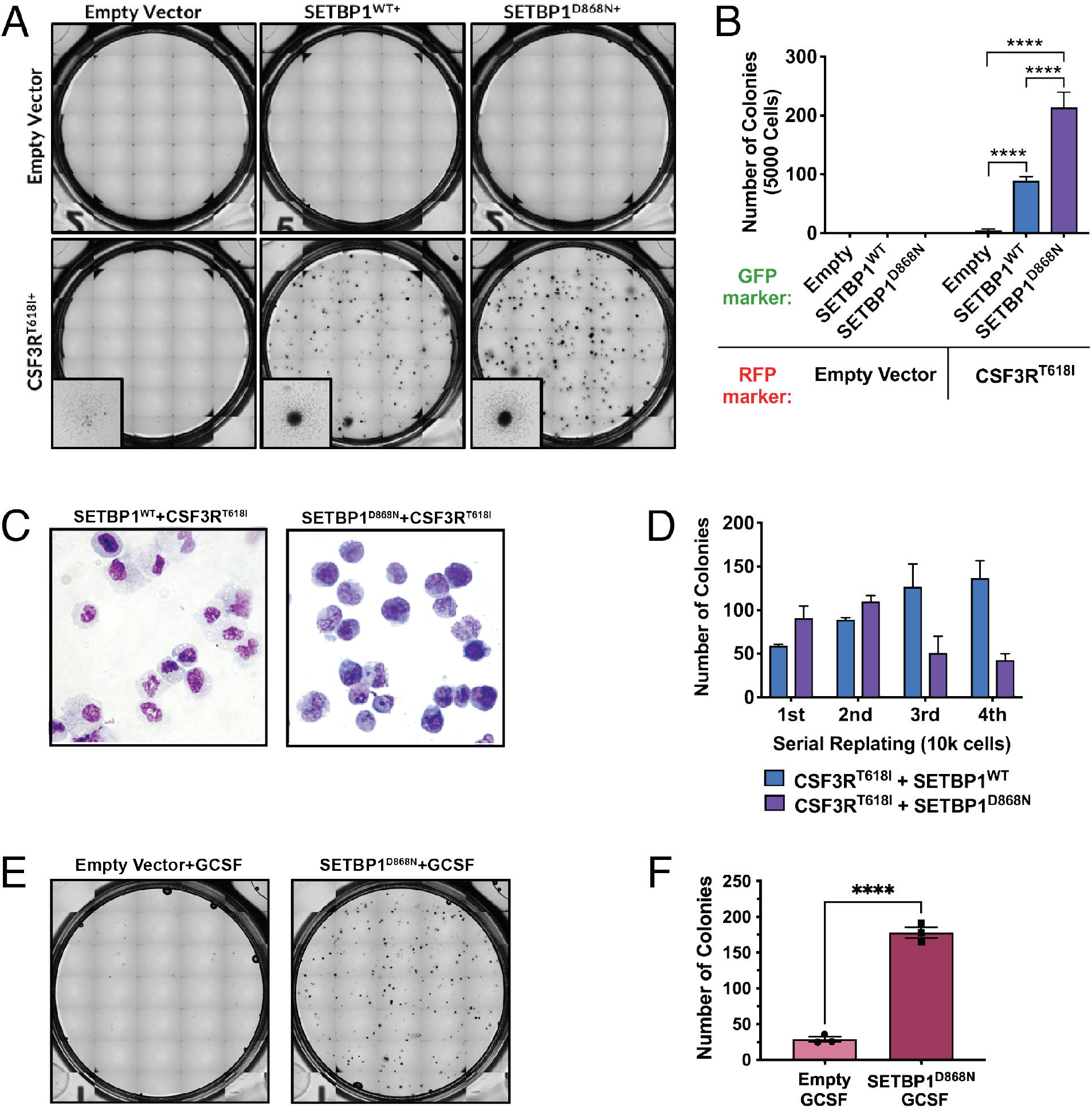
*SETBP1* combines with *CSF3R* mutations to promote cellular proliferation *in vitro*. **(A)** To evaluate the effects of *SETBP1*^*Wt*^, *SETBP1*^*D868N*^, and *CSF3R*^*T618I*^ or the combination of these mutations on hematopoietic progenitors, mouse bone marrow was retrovirally transduced to express mutations of interest or appropriate retroviral control vectors. Cells were then sorted based on fluorescent markers and plated in cytokine-free methylcellulose media in triplicate for a colony forming unit (CFU) assay. Representative images are shown here at day 7. **(B)** Quantification of the colony phenotype shown in A. Statistics: Two-Way ANOVA with Tukey correction, shown for key relationships. Both the *CSF3R*^*T618I*^*+SETBP1*^*Wt*^ and *CSF3R*^*T618I*^*+SETBP1*^*D868N*^ groups are significantly higher than every group with an empty vector (p<0.0001). ^*^p<0.05, ^**^p<0.01, ^***^p<0.001, ^****^p<0.0001 **(C)** Individual colonies were harvested from the methylcellulose using a glass pipette and spread onto a glass slide. Slides were then allowed to dry for 4-6 hours, stained with May-Grünwald and Giemsa solutions and imaged. Representative images shown for cells expressing *CSF3R*^T6181^ with either *SETBP1*^*Wt*^ or *SETBP1*^D868N^. **(D)** After 7 days in culture, cells were harvested by diluting the methylcellulose with PBS and performing three PBS washes. Cells were counted using a TC20 and approximately 10,000 cells per condition were plated into fresh cytokine-free Methylcellulose media in triplicate. Serial replating is successful to at least four passages with both *CSF3R*^*T618I*^+*SETBP1*^*Wt*^ and *CSF3R*^*T618I*^+*SETBP1*^*D868N*^. **(E)** To evaluate if *SETBP1*^D868N^ enhanced colony formation driven by the endogenous ligand for CSF3R (GCSF), we plated 8,000 *SETBP1*^D868N^ expressing cells in cytokine-free methylcellulose media with or without exogenous GCSF (100 ng/ml). Representative images are shown. **(F)** Quantification of the CFU assay in E, with unpaired two-tailed t-test. ^*^p<0.05, ^**^p<0.01, ^***^p<0.001, ^****^p<0.0001

Since transgenic models are not yet available for mutant *SETBP1*, we used retroviral vectors to study whether *SETBP1*^*D868N*^ augments *CSF3R*^*T618I*^-driven oncogenesis *in vivo*. When 25,000 lineage-negative Balb/c bone marrow cells expressing *CSF3R*^*T618I*^ and/or *SETBP1*^*D868N*^ were transplanted into lethally irradiated mice along with 250,000 carrier cells, the mice with both mutations developed an aggressive myeloid leukemia in less than three weeks (**Figure 2A**). This was associated with a rapid expansion of the granulocyte lineage, massive splenomegaly, and moderate hepatomegaly (**Figure 2B-F**). There were no significant changes in terminal body weight (**Figure 2G**). Mice transplanted with bone marrow expressing *SETBP1*^*D868N*^ alone had a median survival of 181 days, while mice receiving *CSF3R*^*T618I*^ alone did not reach their median survival during the course of this study (**Figure 2A**). A second transplant was performed using bone marrow from Balb/c donors that had been treated with Fluorouracil (5FU) to deplete mature progenitor cells. Transduced 5FU-treated marrow was sorted and 2,000 cells per condition— along with 200,000 carrier marrow cells—were transplanted into lethally irradiated mice (**Figure S1A**). At Day 19, mice were sacrificed to collect flow cytometry endpoint on the bone marrow compartment. Mice with *CSF3R*^*T618I*^+*SETBP1*^*D868N*^ marrow had granulocytosis with an expansion of Cd11b+ cells in the blood and bone marrow (**Figure S1B-C**).

**Figure 2.**
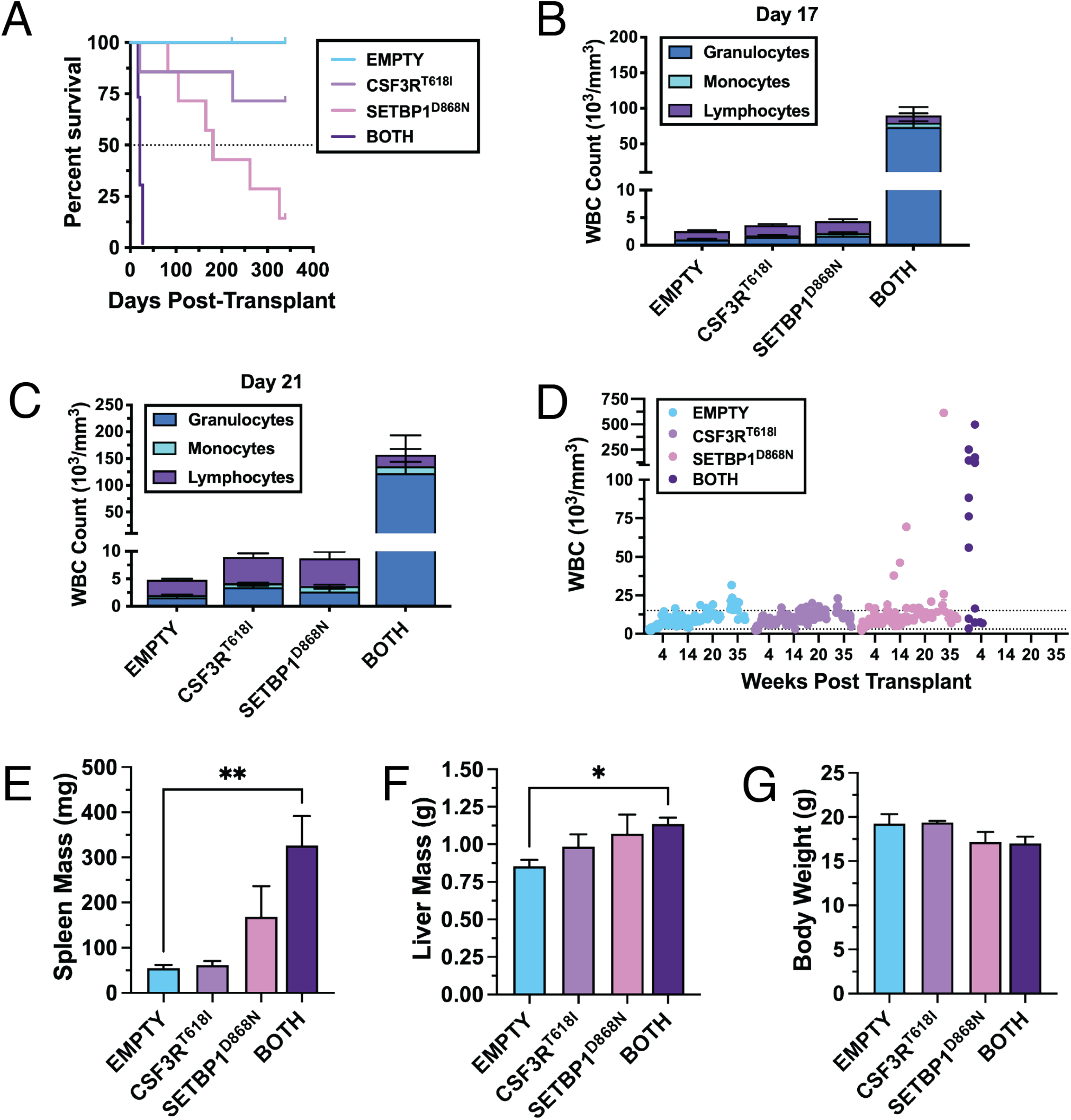
Transplantation of bone marrow cells expressing *CSF3R* and *SETBP1* mutations results in a rapidly lethal leukemia. **(A)** Survival curves for *SETBP1* primary transplant model. Transplantation of 25,000 lineage negative *CSF3R*^*T618I*^+*SETBP1*^*D868N*^ cells—with 190,000 non-transfected carrier bone marrow cells—into lethally irradiated recipient mice resulted in a rapidly lethal, aggressive leukemia. The median survival of *SETBP1*^*D868N*^ mice was 25.9 weeks. **(B)** Peripheral blood white blood cell (WBC) differentials at post-transplant day 17. **(C)** Peripheral WBC differentials at day 21. **(D)** Peripheral WBC counts over time. **(E)** Terminal spleen mass. **(F)** Terminal liver mass. **(G)** Terminal body weights. Statistics: One-Way ANOVA with Dunnett correction. ^*^p<0.05, ^**^p<0.01, ^***^p<0.001, ^****^p<0.0001

Since co-expression of *CSF3R*^*T618I*^ and *SETBP1*^*Wt or D868N*^ conferred replating potential in cytokine-free CFU assays, we hypothesized that these cells might also proliferate in liquid culture. Indeed, we discovered that *CSF3R*^*T618I*^+*SETBP1*^*Wt*^ and *CSF3R*^*T618I*^+*SETBP1*^*D868N*^ expressing cells harvested from CFU assay grow in IMDM with 20% FBS and no cytokine supplementation (**Figure S2A-B**). These cells can be maintained in culture for months with continued cell division and high viability. Neither gene alone confers this growth potential (data not shown). To understand how *SETBP1* expression confers hemopoietic cell expansion in the context of *CSF3R*^*T618I*^, we generated a new cell line in which expression of *SETBP1*^*D868N*^ is regulated by doxycycline (**Figure 3A**). Withdrawal of doxycycline from the cell culture media silences expression of *SETBP1*^*D868N*^and results in a cessation of cell growth after 48 hours and a sharp drop in viability at 72 hours (**Figure 3B,C**). At 24 hours, cells cultured with and without doxycycline have similar Cd11b and GR-1 expression, indicating they are at comparable myeloid differentiation states (**Figure 3D,E**). At 48 hours post-doxycycline withdrawal, there is a significant increase in the percentage of cells with high GR-1 expression and there is a clear morphological difference in the cells by histology (**Figure 3E,F**). Cells expressing only *CSF3R*^*T618I*^ (doxycycline negative) differentiate into mature myeloid cells, including neutrophil precursors and neutrophils with ring-shaped nuclei. This shift in differentiation at 48 hours post-*SETBP1*^*D868N*^ silencing is accompanied by cell cycle arrest in G0/G1 (**Figure 3G,H**) and is consistent with proliferation and differentiation acting in opposition. This *CSF3R*^*T618I*^*+SETBP1*^*D868N*-dox^ cell line model provides a tractable system in which to evaluate SETBP1-driven molecular programs.

**Figure 3.**
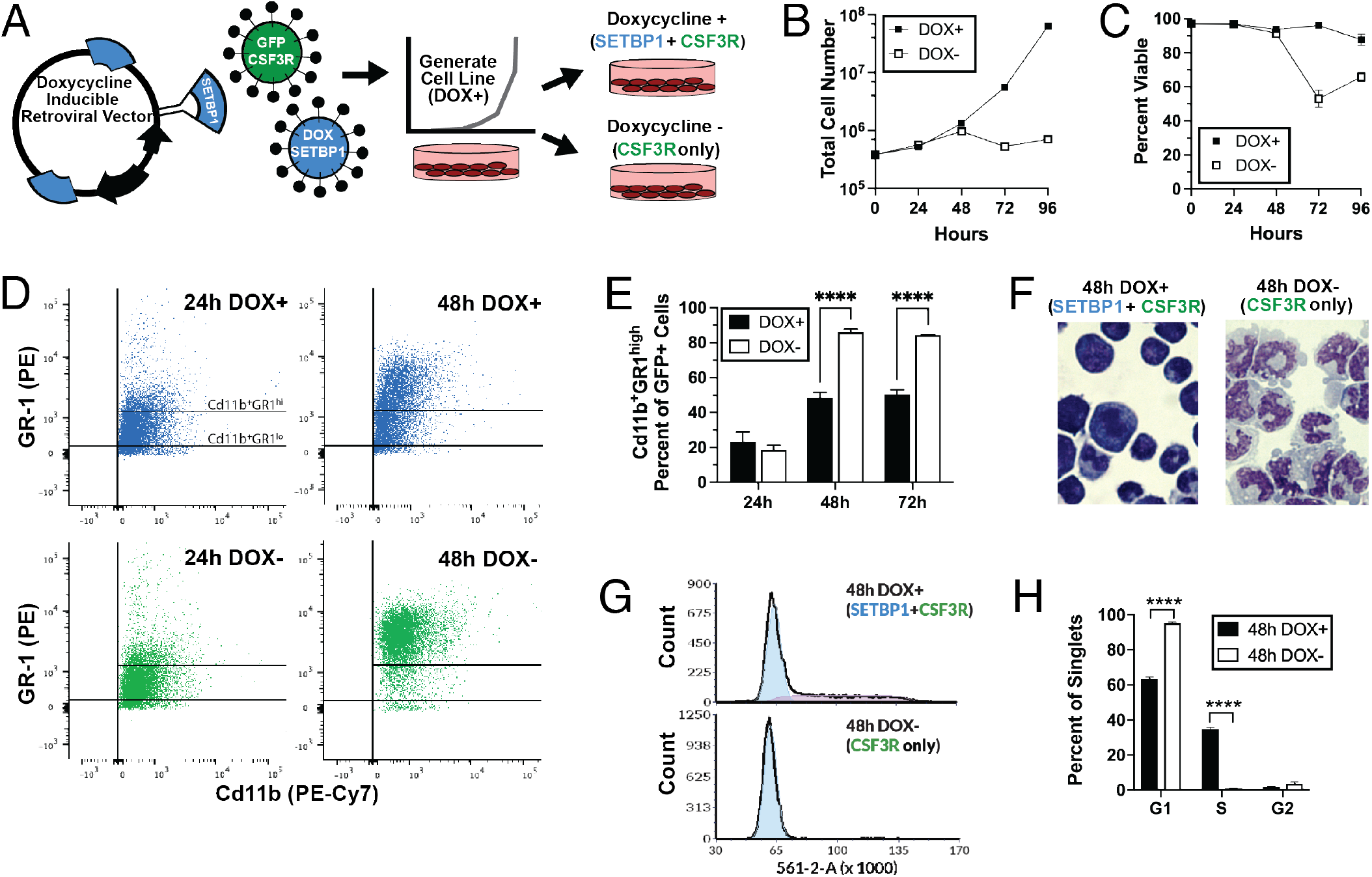
*CSF3R*^*T618I*^ and *SETBP1*^D868N^expressing hematopoietic cells undergo cell cycle arrest and differentiation following SETBP1 withdrawal. **(A)** Schematic of doxycycline-inducible (DOX) cell line generation (*CSF3R*^*T618I*^*+SETBP1*^*D868N*-dox^). This cell line was generated from primary mouse bone marrow using retrovirally expressed oncogenes, with *CSF3R*^*T618I*^ in a constitutively active vector and *SETBP1*^*D868N*^ in a Tet-on vector. Following transduction with both oncogenes, *CSF3R*+ cells (GFP+) were sorted and then cultured in the presence of doxycycline (1ug/ml) to induce *SETBP1*^*D868N*^ expression. **(B)** Growth of *CSF3R*^*T618I*^*+SETBP1*^*D868N*-dox^ cells with and without doxycycline. To shut off *SETBP1* expression, cells were washed with PBS five times and then resuspended in media +/- doxycycline in triplicate. Cells expressing only *CSF3R*^*T618I*^ stop proliferating after 48 hours. **(C)** Cell death increases between 48 and 72 hours following *SETBP1*-withdrawal. **(D)** Representative flow cytometry plots for Cd11b and GR1 expression at 24 and 48 hours post withdrawal. After withdrawing doxycycline, cells were collected at 24-hour intervals to monitor changes in cell state. **(E)** Quantification of Cd11b^+^GR1^high^ cells with and without doxycycline at 24, 48 and 72 hours. Statistics: Repeated measures ANOVA with multiple comparisons within timepoints, Šidák correction. ^*^p<0.05, ^**^p<0.01, ^***^p<0.001, ^****^p<0.0001 **(F)** Representative images of the cell line undergoing differentiation at 48 hours. **(G)** At 48 hours post-withdrawal, cells without *SETBP1* expression are arrested in G0/G1 as measured by flow cytometry using propidium iodide staining on fixed cells. **(H)** Quantification of cell cycle phases from G at 48 hours. Statistics: Two-Way ANOVA with significance for comparisons within cell cycle phase shown, Šidák correction. ^*^p<0.05, ^**^p<0.01, ^***^p<0.001, ^****^p<0.0001

To identify transcriptional programs that are upregulated by *SETBP1*^*D868N*^ in the context of CNL, we performed RNA-seq *CSF3R*^*T618I*^*+SETBP1*^*D868N*-dox^ cell line at 24 hours post-doxycycline-withdrawal, when the cells were still viable and dividing, and compared them to cells treated with doxycycline. One of the strongest signatures in cells expressing *SETBP1*^*D868N*^ relative to those without *SETBP1*^*D868N*^ is that of MYC target genes (**Figure 4A-D)**. Pathway analysis of the differentially expressed genes between *CSF3R*^*T618I*^ only (doxycycline negative) and *CSF3R*^*T618I*^+*SETBP1*^*D868N*^ (doxycycline positive) showed that pathways up with *SETBP1*^*D868N*^ were overwhelmingly associated with MYC perturbations (**Figure 4A**). In the *CSF3R*^*T618I*^-only condition, Brown Myeloid Cell Development differentiation-associated genes were enriched (**Figure 4B**). This is in line with our data showing that the *CSF3R*^*T618I*^-only cells differentiate into mature myeloid cells between 24 and 48h post-doxycycline withdrawal (**Figure 3D-F**). Congruent with previous studies of SETBP1^7-9,11^, Gene Set Enrichment Analysis (GSEA) showed that SETBP1-associated genes are enriched for early progenitor pathways, including upregulated targets of Hoxa9 and Meis1 (**Figure 4B**). Consistent with the pathway analysis in **Figure 4A**, GSEA also identified that MYC targets were associated with *SETBP1*^*D868N*^ (**Figure 4B**). At the individual gene level, we find that *Myc, Meis1* and *Hoxa9* themselves highly upregulated (**Figure 4C**). Additionally, we see that *Hoxa10* and *Myb*, which have been previously associated with *SETBP1-*driven leukemogenesis^7,11^, are among the top differentially regulated genes when *SETBP1* is expressed (**Figure 4C**).

**Figure 4.**
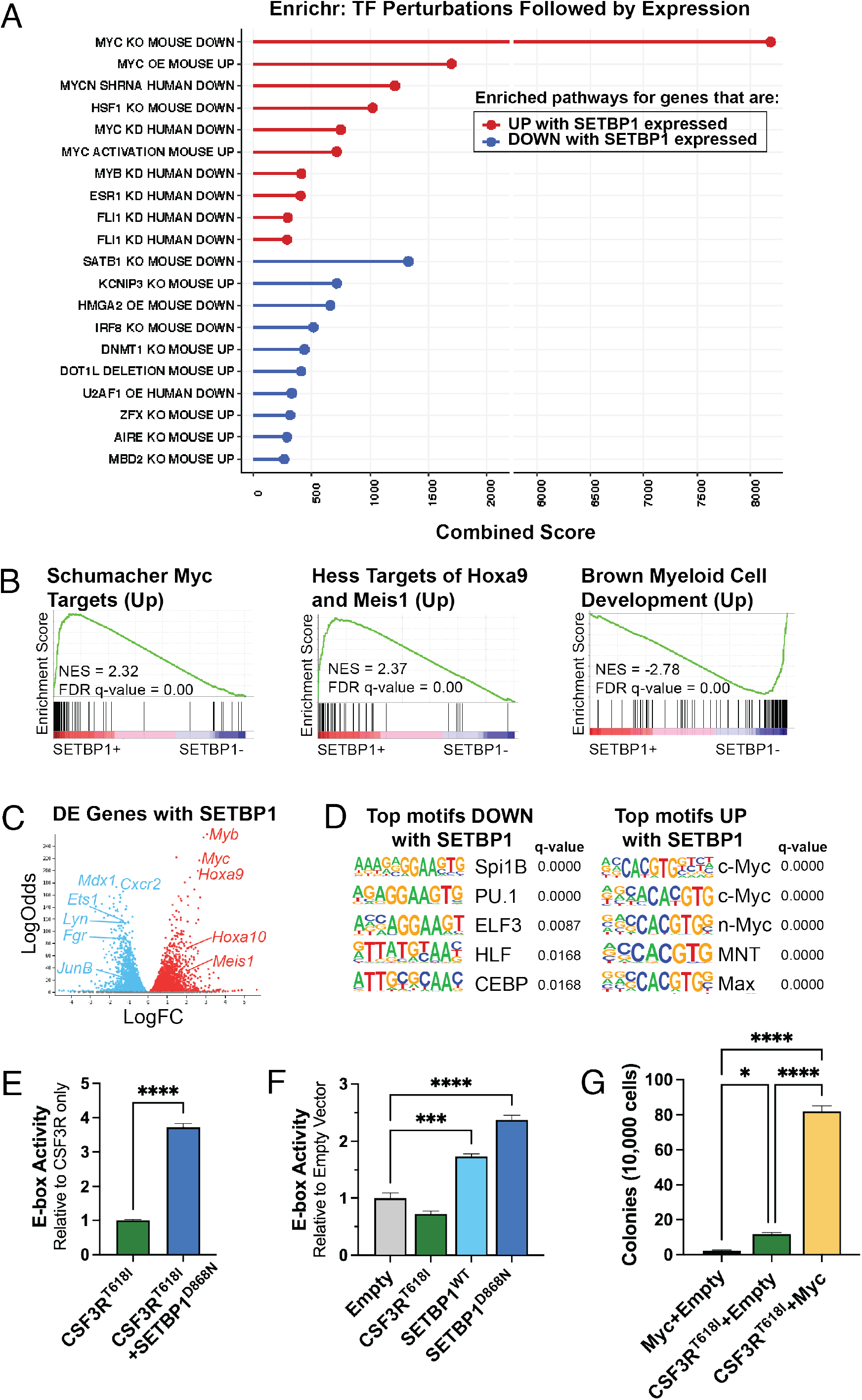
SETBP1 upregulates early progenitor gene expression pathways and is associated with increased activation of MYC targets. **(A)** Top Enrichr “transcription factor perturbation followed by expression” pathways for genes that are differentially expressed with *SETBP1*. We performed RNA-seq on the *CSF3R*^*T618I*^*+SETBP1*^*D868N*-dox^ cell line at 24 hours post-doxycycline-withdrawal, when the cells were still viable and dividing. Data are reported for cells with doxycycline (*CSF3R*^*T618I*^*+SETBP1*^*D868N-ON*^) relative to without doxycycline (*CSF3R*^*T618I*^*+SETBP1*^*D868N-OFF*^). Enrichr calculates the combined score by multiplying the pathway z-score and log(p-value). **(B)** Gene set enrichment analysis (GSEA) was performed to identify pathways that are enriched with *SETBP1* expressed. Three of the top GSEA plots, each with an FDR q-value of 0.00, are shown. **(C)** Glimma volcano plot showing differentially expressed genes with SETBP1, with several key genes annotated. **(D)** HOMER motif analysis was run to identify the top motifs enriched in the genes that are differentially upregulated and downregulated by SETBP1. **(E)** Co-expressing *SETBP1*^*D868N*^ with *CSF3R*^*T618I*^ drives a 3.7-fold increase in MYC activity over *CSF3R*^*T618I*^ alone. A luciferase reporter assay for the MYC E-box was utilized to measure if *SETBP1*^*D868N*^ modulates E-box activity. Using a MYC E-box reporter plasmid, E-box activity was measured in transfected 293T17 cells expressing *CSF3R*^*T618I*^ alone or *CSF3R*^*T618I*^+SETBP1^D868N^. **(F)** In transfected 293T17 cells expressing only *CSF3R*^*T618I*^, *SETBP1*^*Wt*^ or *SETBP1*^*D868N*^, *CSF3R* does not increase E-box activity above baseline while both wildtype and mutant SETBP1 significantly increase E-box activity. **(G)** Co-expressing *MYC* with *CSF3R*^*T618I*^ in a CFU assay results in an increase in colony formation over either oncogene alone. A colony forming unit assay was performed to assess whether expression of *MYC* is sufficient to increase *CSF3R*-driven colony formation. 10,000 cells expressing either *MYC, CSF3R*^T6181^ or both were plated in cytokine-free methylcellulose and CFUs were counted after 7 days. ^*^p<0.05, ^**^p<0.01, ^***^p<0.001, ^****^p<0.0001

To determine if there were particular motifs enriched in regulatory regions of differentially expressed genes, we ran the HOMER motif discovery algorithm^13^. This analysis identified that the genes upregulated by *SETBP1*^*D868N*^ are enriched for genes regulated by MYC E-box motifs (**Figure 4D**). To validate this finding, we used a luciferase reporter driven by the MYC E-box element to measure if *SETBP1*^*D868N*^ modulates E-box activity. Congruent with the RNA-seq analysis, co-expressing *SETBP1*^*D868N*^ with *CSF3R*^*T618I*^ drives a 3.7-fold increase in MYC activity over *CSF3R*^*T618I*^ alone (**Figure 4E**). Independent of *CSF3R*^*T618I*^, both *SETBP1*^*Wt*^ and *SETBP1*^*D868N*^ increase MYC E-box activity by 1.7-fold and 2.4-fold, respectively (**Figure 4F**). Using a CFU assay, we demonstrate the retroviral overexpression of *MYC* is sufficient to enhance *CSF3R*^*T618I*^-driven colony formation (**Figure 4G**).

To better understand the epigenetic changes associated with these differential gene expression programs, we performed CUT&Tag in the *CSF3R*^*T618I*^*+SETBP1*^*D868N*-dox^ cell line for three histone marks: H3K4me1, H3K4me3 and H3K27Ac (**Figure 5A**). H3K4me is primarily associated with enhancers and H3K4me3 with promoters. H3K27Ac is associated with both active promoters and active enhancers. While there was not a global change in deposition of these epigenetic marks, H3K4me3 and H3K27Ac differential peaks have enhanced MYC/MYB motif enrichment when *SETBP1*^*D868N*^ is expressed (**Figure S3A, Figure 5B**). Congruent with the RNA-seq data, MYC motifs were enriched in the peaks that are upregulated by SETBP1 (**Figure 5B, Figure S3B**).

**Figure 5.**
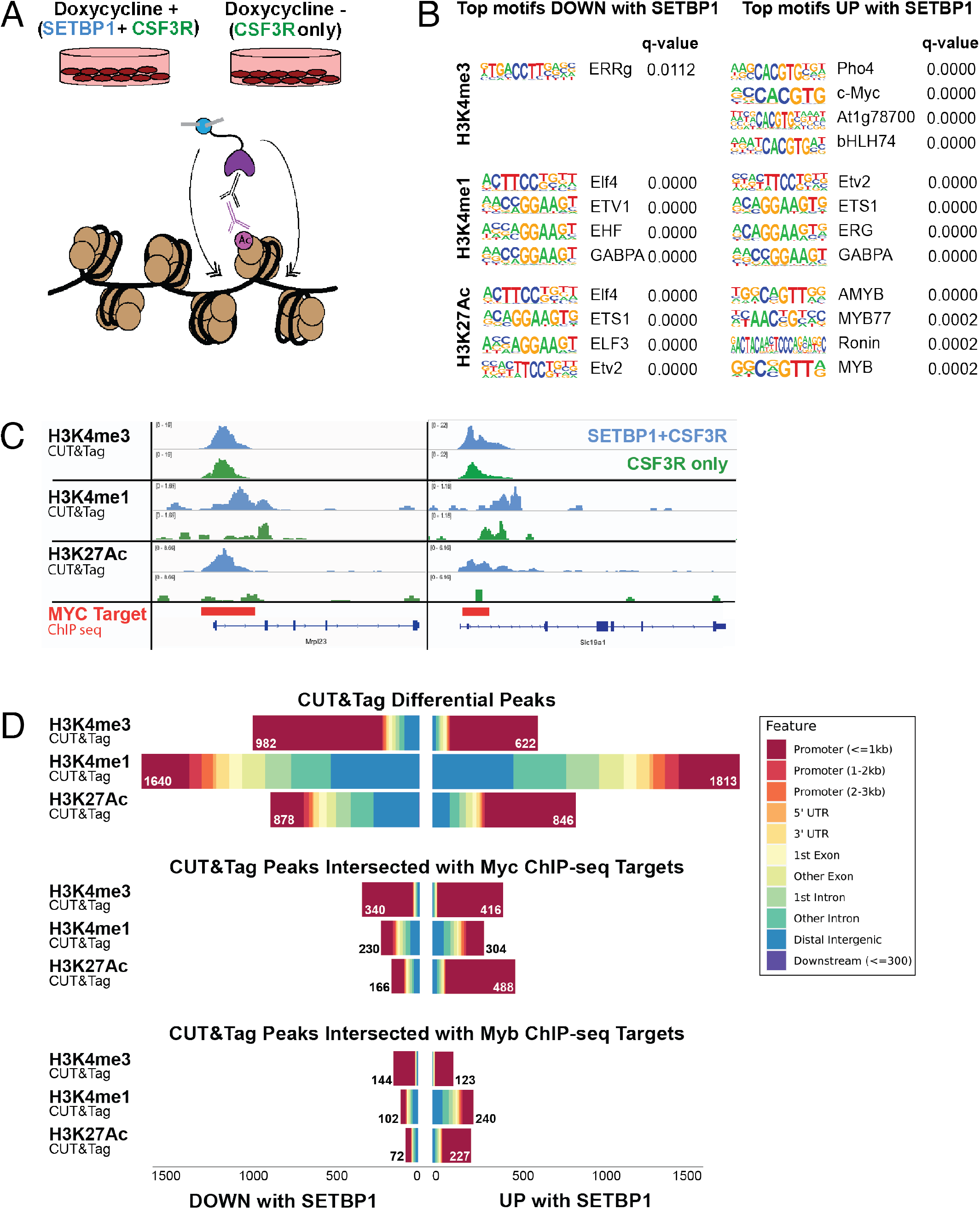
Epigenetic regulation of MYC targets by SETBP1. **(A)** Schematic: we performed CUT&Tag in our *CSF3R*^*T618I*^*+SETBP1*^*D868N*-dox^ cell line for three histone marks. Doxycycline was withdrawn from the cell line to turn off oncogenic SETBP1 expression and cells were harvested for epigenetic analyses at 24 hours post-withdrawal. **(B)** Top four motifs identified by HOMER. Enriched motifs in each histone mark were determined using the genomic coordinates of differential peaks. Motifs UP with SETBP1 are enriched when SETBP1 is expressed (DOX+) relative to without SETBP1 (DOX-). **(C)** Representative tracks are shown here for each histone mark at the location of a MYC target identified using MYC ChIP-seq data from ENCODE (ENCSR000EUA). **(D)** Features plots for differential peaks, showing the breakdown of peaks within promoters and other elements. Regions of significant SETBP1-induced histone peaks were intersected with either MYC (ENCFF152JNC) or MYB (ENCFF911NHJ) ChIP-seq data. The total number of differential peaks for each condition is annotated.

Using a public Myc ChIP-seq dataset (ENCFF152JNC), we intersected intervals of Myc binding with regions of significant SETBP1-induced peaks. Remarkably, 47% of the differential H3K4me3 peaks overlapped with Myc binding regions (756/1,604 peaks), indicating an overlap in the promoters that are differentially regulated by Myc and SETBP1^D868N^ (**Figure 5C,D**). The overlap between differential H3K4me1 peaks and Myc targets was 15% (534/3,453), and there was 38% overlap for H3K27Ac and Myc (654/1,724). For Myb bound regions (ENCFF911NHJ), there were fewer regions of overlap: H3K4me3 (17%: 267/1,604 peaks), H3K4me1 (9%: 308/3,453), H3K27Ac (17%: 299/1,724) (**Figure 5D**). We next set out to determine whether the aberrant programs might be pharmacologically reversible.

To determine the essential cell growth and survival pathways in *SETBP1* mutated cells, we performed a chemical screen with 175 inhibitors with known sensitivity in patient samples (BeatAML cohort^14^). The median IC50 for each inhibitor in the BeatAML cohort^14^ was divided by the IC50 for the same inhibitor in the *CSF3R*^*T618I*^+*SETBP1*^*D868N*^ cell line (two biological replicate lines, technical triplicates) to calculate a fold increase in sensitivity relative to other samples. This enabled us to look at which drugs this sample is particularly sensitive to, as opposed to drugs that are generally toxic. Consistent with the activation of the JAK/STAT pathway by mutant *CSF3R*, this cell line was sensitive to JAK inhibitors. Interestingly, the top two hits were lysine specific demethylase 1 (LSD1) inhibitors (**Figure 6A, Figure S4A**). Annotations for the top 15 inhibitors are found in **Table S1** and **Table S2**. A previous study found that LSD1 induces *Myc* expression in a non-hematopoietic context^15^. We were therefore interested in whether LSD1 inhibitors can reduce aberrant MYC activity driven by SETBP1. Using a luciferase promoter assay, we determined that MYC E-box activity was modulated by LSD1 inhibition and found a modest dose-dependent response to GSK2879552, culminating in a 24% reduction in E-box activity at 250 nM (**Figure 6B**). We next tested a third LSD1 inhibitor, GSK-LSD1, which proved to be more potent in this cell line with an IC50 of approximately 250 nM compared to 590 nM (**Figure S4B**).

**Figure 6.**
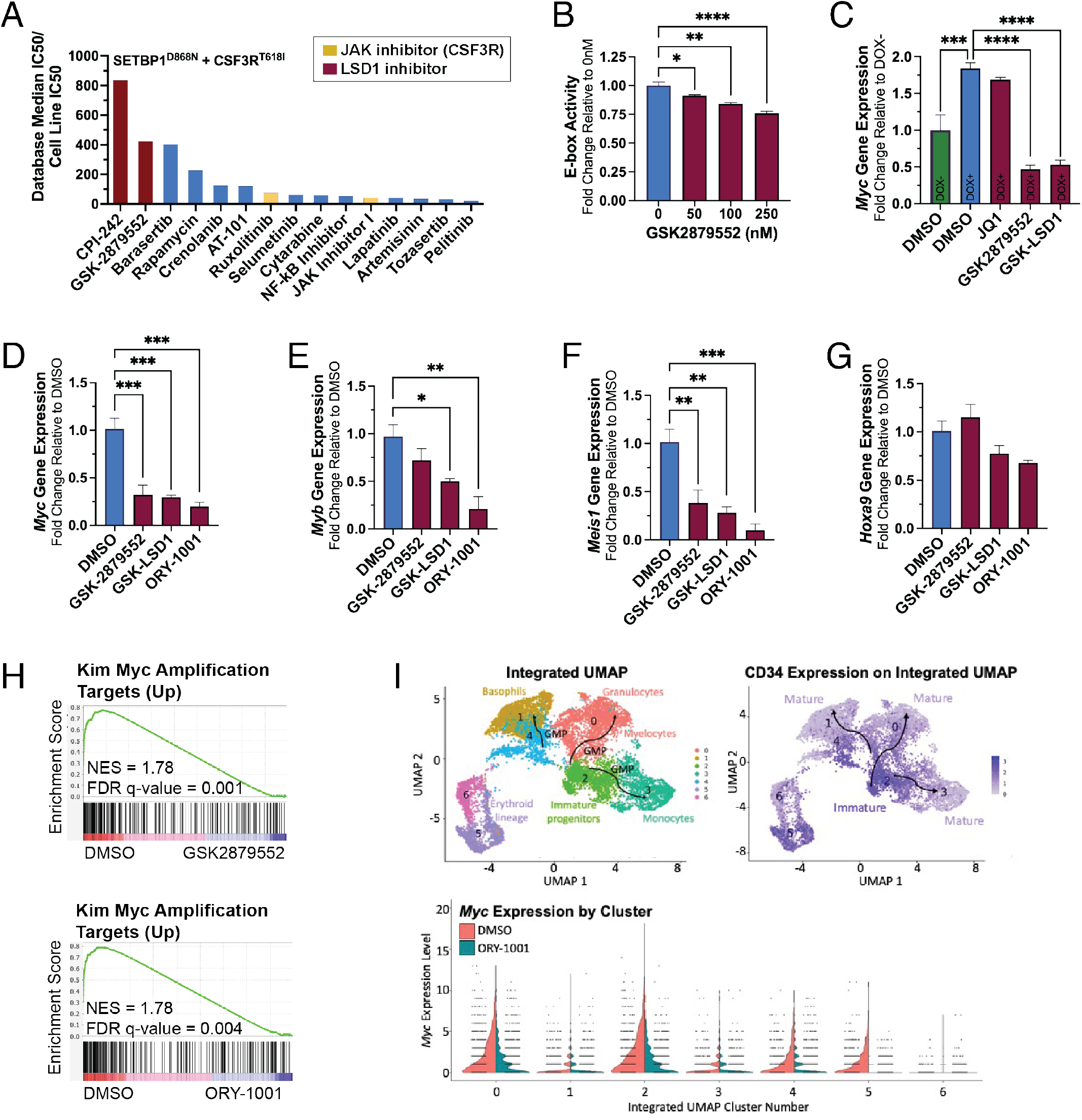
LSD1 inhibitors normalize aberrant SETBP1 transcriptional programs. **(A)** A medium throughput inhibitor screen was performed on the *CSF3R*^*T618I*^+*SETBP1*^*D868N*^ cell line and both LSD1 inhibitors and JAK inhibitors were among the top hits. The 175 inhibitors evaluated have known sensitivity in patient samples (BeatAML cohort^14^). The inhibitors were ranked for this analysis by dividing the median IC50 of all samples previously screened by our cell line IC50 to determine a fold change. **(B)** A luciferase E-box activity assay was performed with four concentrations of the LSD1 inhibitor GSK2879552. In 293T17 cells expressing *CSF3R*^*T618I*^ and *SETBP1*^*D868N*^, LSD1 inhibition reduced MYC activity by 24% at 250 nM. **(C)** In our cell line where *SETBP1*^*D868N*^ expression is regulated by doxycycline (DOX), we evaluated whether LSD1 inhibitors would reduce *Myc* gene expression to the level of DOX-cells. The LSD1 inhibitors GSK2879552 (1000nM) and GSK-LSD1 (100nM) both reduce *Myc* expression in *CSF3R*^*T618I*^+*SETBP1*^*D868N*^ cells, but not JQ1 (200nM). **(D)** qPCR for *Myc* was performed following treatment of the *CSF3R*^*T618I*^+*SETBP1*^*D868N*^ cell line with one of three LSD1 inhibitors at 100 nM (GSK2879552) or 30 nM (GSK-LSD1, ORY-1001) for 48 hours. **(E)** qPCR for *Myb*. **(F)** qPCR for *Meis1* **(G)** qPCR for *Hoxa9*, which is not modulated by LSD1 inhibition at these concentrations. **(H)** RNA-seq was performed following treatment of the cell line with 100 nM GSK2879552 or 30 nM ORY-1001 for 24 hours. GSEA demonstrates that this treatment is associated with a reversal of MYC amplification with both inhibitors. **(I)** A *CSF3R*^*T618I*^ and *SETBP1*^*G870S*^ mutated patient sample was treated with 100 nM ORY-1001 for 24h and CITE-seq (single cell RNA-seq with barcoded antibody labeling) was performed. Treatment significantly decreases *Myc* expression in hematopoetic progentior clusters expressing high levels of CD34. ^*^p<0.05, ^**^p<0.01, ^***^p<0.001, ^****^p<0.0001

While the *CSF3R*^*T618I*^*+SETBP1*^*D868N*^ cell line was sensitive to the LSD1 inhibitors, cell death occurred at higher doses (>100 nM). Given this, we were interested in understanding whether LSD1 treatment could modulate SETBP1-driven oncogenic programs and if lower doses could potentially sensitize cells to other therapies. To determine whether LSD1 inhibition reduces *Myc* gene expression to basal levels, we utilized the *CSF3R*^*T618I*^*+SETBP1*^*D868N*-dox^ cell line. To determine basal gene expression, *SETBP1* ^*D868N*^ expression was silenced by withdrawing doxycycline in triplicate. In parallel, *CSF3R*^*T618I*^*+SETBP1*^*D868N*-dox^ cells were cultured in the presence of doxycycline and treated in triplicate with either DMSO, JQ1, GSK2879552 or GSK-LSD1. After 48 hours, cells were harvested to assess *Myc* expression by qPCR. Treatment with the bromodomain inhibitor JQ1, which targets *Myc*, yielded no significant changes to *Myc* expression in this molecular context. However, both LSD1 inhibitors reduced *Myc* expression significantly (**Figure 6C**). Using the cell line that constitutively expresses *CSF3R*^*T618I*^*+SETBP1*^*D868N*^, we then evaluated how lower doses of LSD1 inhibition (100 nM GSK2879552, 30 nM GSK-LSD1 or 30 nM ORY-1001) modulated four key SETBP1-associated genes: *Myc, Myb, Meis1*, and *Hoxa9*. In the *CSF3R*^*T618I*^*+SETBP1*^*D868N*^ cells, LSD1 inhibitors reduced *Myc, Myb* and *Meis1* expression, but did not significantly decrease *Hoxa9* expression after 48 hours of treatment (**Figure 6D-G**). Of note, another inhibitor of LSD1 under investigation in clinical trials—ORY-1001—was remarkably effective, reducing *Myc, Myb* and *Meis1 expression* by approximately 80-90%. To assess global transcriptional changes with LSD1 inhibition, we performed RNA-seq on cells treated with either 100 nM GSK2879552 or 30 nM ORY-1001 for 24 hours. Using GSEA, we see that LSD1 inhibition is inversely associated with MYC target amplification—MYC targets are enriched in the DMSO treated cells relative to the LSD1 treated cells (**Figure 6H**).

CNL patient samples are rare and can exhibit low viability following cryopreservation due to the abundance of neutrophils in the peripheral blood and bone marrow. To assess whether LSD1 inhibition could modulate progenitor populations and MYC signaling in a human patient sample, we isolated viable CD34+ progenitor cells from a *CSF3R*^*T816I*^+*SETBP1*^*G870S*^ cryopreserved CNL bone marrow sample and cultured these CD34+ cells in a serum-free expansion media for 7 days. The total number of CD34+ cells expanded from 65,400 to 642,000 cells in 7 days. 300,000 cells were then treated with either 100 nM ORY-1001 or DMSO for 24 hours. After treatment, single cell RNA-seq with barcoded antibody labeling (CITE-seq) was performed. Marker genes (*MPO, GATA1, GATA2, IRF8, ELANE, LYZ, CEBPE*) and surface antigens (CD34, CD45RA) were used for population identification (**Figure 6I, Figure S5**). We found that ORY-1001 treatment significantly decreases *MYC* expression in hematopoetic progentior clusters expressing high levels of CD34 (**Figure 6I**).

The JAK inhibitor ruxolitinib is under investigation as a promising therapeutic agent for patients who have mutations in *CSF3R*, and has shown efficacy in clinical trial^2^. To improve initial treatment response rates and circumvent resistance, it is likely that a multipronged therapeutic approach will be needed. From our chemical screen in the *CSF3R*^*T618I*^*+SETBP1*^*D868N*^ cell line, we knew that these cells are sensitive to JAK inhibitors relative to the median IC50 for patient samples in the BeatAML cohort^14^ (**Figure 6A**). To evaluate how *SETBP1*^*D868N*^ alters sensitivity to ruxolitinib, we performed a 7-day cytokine-free colony assay with mouse bone marrow retrovirally transduced with either *CSF3R*^T6181^+empty vector or *CSF3R*^T6181^+*SETBP1*^D868N^ (**Figure S6A**). Cells were plated with increasing concentrations of ruxolitinib and found to have less sensitivity with *CSF3R*^T6181^+*SETBP1*^D868N^ (IC50 296 nM) than with *CSF3R*^T6181^+empty vector (78 nM). The IC50 of primary *CSF3R*^T6181^+*SETBP1*^D868N^ transduced cells in colony assay is similar to the *CSF3R*^*T618I*^*+SETBP1*^*D868N*^ cell line (241 nM) (**Figure S6B**).

To determine whether LSD1 inhibitors, which reduce SETBP1-associated aberrant gene expression, are effective in combination with JAK inhibitors, we next evaluated the synergy between these two agents (**Figure 7A, Figure S6C**). Each LSD1 inhibitor that we tested exhibited marked synergy with ruxolitinib, with the greatest synergy observed with ORY-1001 (delta score 22.028). To understand the mechanisms underlying this drug synergy, RNA-seq was performed on cells treated with DMSO, ruxolinitib, GSK2879552, ORY-1001, ruxolitinib with GSK2879552 or ruxolitinib with ORY-1001. A heatmap was generated using unbiased clustering and the individual clusters were analyzed using HOMER motif enrichment and Enrichr pathway analysis (**Figure 7B**). In Cluster 1, we see genes that are upregulated by the combination therapy more than either drug alone. Motifs for Cluster 1 include differentiation-associated transcription factors PU.1 and Runx1, which are both members of the core binding factor (CBF) complex. Cluster 4 contains genes that are downregulated more by the combination than either drug alone. In this cluster, we see a Myc and Fli1 signature. This supports a model where LSD1 inhibition is reversing SETBP1-associated phenotypes, and provides rationale for combined therapeutic strategies utilizing these epigenetic regulators in combination with ruxolitinib leukemia with both *CSF3R* and *SETBP1* mutations.

**Figure 7.**
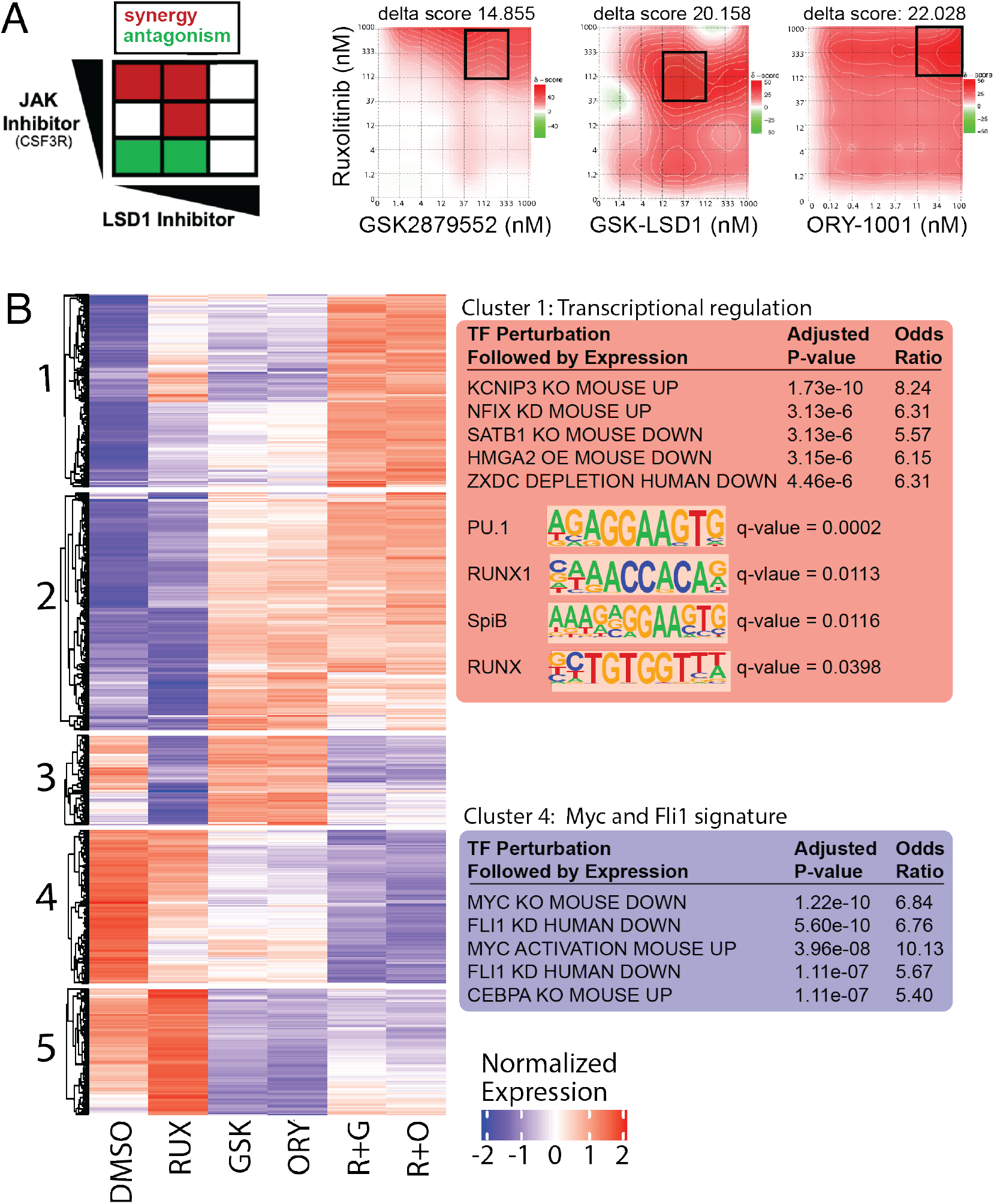
**(A)** Synergy between each LSD1 inhibitor and ruxolitinib, with the greatest synergy between ORY-1001 and ruxolitinib (delta score: 22.028). To determine if there is synergy between LSD1 inhibition targeting SETBP1-driven pathways and a JAK inhibitor (ruxolitinib) targeting CSF3R-driven pathways, the *CSF3R*^*T618I*^+*SETBP1*^*D868N*^ cell line was plated in an 8×8 matrix in triplicate with increasing concentrations of each inhibitor. **(B)** RNA-seq data from cells treated with DMSO, 100 nM ruxolinitib, 100 nM GSK2879552 (GSK), 30 nM ORY-1001, ruxolitinib with GSK2879552 (R+G) or ruxolitinib with ORY-1001 (R+O). Cluster 1 represents pathways upregulated more by the combination therapy than either drug alone and includes a number of differentiation-associated transcription factors. Cluster 4 represents pathways downregulated more by the combination than either drug alone and has a Myc and Fli1 signature.

## Discussion

*SETBP1* is recurrently mutated in myeloid malignancies including atypical CML, chronic myelomonocytic leukemia (CMML) and juvenile myelomonocytic leukemia (JMML). *SETBP1* mutations are at a particularly high frequency in CNL—a leukemia characterized by *CSF3R* mutations and the overproduction of neutrophils. The primary goal of this study was to determine how *SETBP1* mutations contribute to the pathobiology of *CSF3R*-driven leukemias. We found that *SETBP1*^*Wt or D868N*^ dramatically increases *CSF3R*^*T618I*^-driven hematopoietic progenitor proliferation and accelerates *CSF3R*^*T618I*^-driven disease (**Figures 1-3**). Expression of *SETBP1*^*D868N*^ leads to upregulation of progenitor-associated gene expression programs and downregulation of differentiation-associated genes. *SETBP1*^*D868N*^-driven Myc expression can be reversed by treatment of cells with LSD1 inhibitors. Furthermore, LSD1-inhibition synergizes with inhibition of *CSF3R*^*T618I*^-driven signaling in these models.

In the context of a *CSF3R* mutation, we found that mutant *SETBP1* increases *Hoxa9, Hoxa10, Meis1* and *Myb* transcript levels and increases their associated gene expression programs (**Figure 4**). This is congruent with previous studies establishing Setbp1 as a transcriptional regulator of Hox genes^7,11^ and *Myb*^11^. An exciting finding of our transcriptional and epigenetic analyses was that expression of *SETBP1*^*D868N*^ is also associated with a strong Myc pathway signature (**Figure 4,5**). Myc is a transcription factor that plays an integral role in establishing a balance between self-renewal and hematopoietic differentiation^17^. The differentiation block and increased proliferation that occurs with *SETBP1*^*D868N*^ in the *CSF3R*^*T618I*^*+SETBP1*^*D868N*-dox^ model is consistent with the known role of Myc in the inhibition of differentiation other leukemia models^17^. Co-expressing *MYC* with *CSF3R*^*T618I*^ in a CFU assay largely recapitulated the dense and proliferative colony phenotype associated with the combination of *CSF3R*^*T618I*^ and *SETBP1*^*D868N*^ (**Figure 4**). This demonstrates that *MYC* overexpression is sufficient for recapitulation of the proliferative phenotype associated with *SETBP1* mutations.

Although we know that ruxolitinib can be efficacious in CNL, it is likely that additional agents will be needed to achieve long term remissions^2^. In our study, we find that while cells with both *CSF3R* and *SETBP1* mutations retain sensitivity to ruxolitinib, cells expressing both mutations have less sensitivity than those with *CSF3R*^*T618I*^ alone (**Figure S6**). We hypothesized that if *SETBP1*^*D868N*^ drives aggressive disease biology through epigenetic dysregulation of Myc regulatory elements, then therapeutic strategies that normalize *Myc* expression would be effective against *SETBP1*-mutant leukemia. We found that LSD1 inhibition decreases cell viability and suppresses aberrant *Myc* expression (**Figure 6**). In a *SETBP1*-mutant CNL patient sample, LSD1 inhibitor treatment significantly decreased *Myc* expression in CD34-high hematopoetic progenitor clusters. Having established that LSD1 inhibition reduces *SETBP1*-driven Myc expression, we next tested whether it might be useful in combination with the JAK inhibition. In this study, each of the three LSD1 inhibitors demonstrated synergy with ruxolitinib and resulted in the repression of Myc pathways (**Figure 7**). RNA-seq analysis revealed that this drug synergy was associated with the reactivation of differentiation associated pathways. A previous study of *CSF3R/CEBPA* mutant AML demonstrated that LSD1 inhibition caused marked reactivation of differentiation-associated enhancers^18^. In our model, genes upregulate by the combination of LSD1 and JAK inhibition were enriched for PU.1 and Runx1 motifs (**Figure 7**). These findings are in line with previous studies of LSD1 inhibition in leukemia showing activation of PU.1 targets in MLL-rearranged AML^19,20^ and KIT-mutant AML^21^.

In summary, we investigated the role of *SETBP1* mutations in *CSF3R* driven-leukemia. We found that *SETBP1* mutations accelerate leukemic progression in mice. When a *SETBP1* mutation is expressed in murine hematopoietic cells along with a *CSF3R* mutation, *SETBP1* promotes proliferation of immature granulocytes through upregulation of the Myc pathway and epigenetic modulation of Myc target genes. Treatment of *CSF3*R and *SETBP1* mutant cells with ruxolitinib (targeting *CSF3R* signaling) and an epigenetic modulatory drug resulted in synergistic cell death and a repression of aberrant transcriptional programs. These data contribute to our understanding of how *SETBP1* mutations augment *CSF3R*-driven oncogenic programs to produce lethal disease, and provide preclinical evidence for a combination therapeutic strategy in *CSF3R* and *SETBP1* mutant leukemia.

## Materials and Methods

### Mice

C57BL/6J mice (Cat#000664) and Balb/cJ mice (#000651) were obtained from The Jackson Laboratories. Experiments were initiated when mice were used between 6-10 weeks of age. Female C57BL/6J mice were used for CFU assays and cell line generation. Female Balb/cJ mice were used for the transplants. All experiments were conducted in accordance with the National Institutes of Health Guide for the Care and Use of Laboratory Animals and approved by the Institutional Animal Care and Use Committee of Oregon Health & Science University (Protocol #TR01_IP00000482). Peripheral blood collected from the saphenous vein was monitored weekly for white blood cell counts (WBC) using a Vet ABC animal blood counter (Scil animal care company). The health of the mice was monitored daily by animal care personnel who were blinded to study groups. Animals were euthanized if WBC reached above 100,000/mm^3^. For survival analysis, a logrank Mantel-Cox test was performed with Bonferroni posthoc correction for multiple comparison.

### Selection of genetic variants

Patients with *SETBP1* mutations most commonly have somatic variants within a four amino acid segment of the SKI-homology domain^1^. These mutations increase the half-life of SETBP1 by perturbing the β-TrCP degron motif, such that E3 ubiquitin ligases cannot recognize the domain to degrade the protein. *SETBP1*^D868N^ is a representative β-TrCP degron motif mutation that co-occurs with *CSF3R* mutations. CNL patients with *CSF3R* mutations have two primary types of mutations: (1) point mutations in the extracellular or transmembrane domain of *CSF3R*, of which the most common is T618I or (2) less common truncating mutations in the cytoplasmic domain^1,22,23^.

### Cloning

For these studies, we utilized V5-tagged-*SETBP1* constructs with codon-optimization, described previously^12^. *SETBP1* constructs were cloned into pMSCV-IRES-mCherry (a gift from Dario Vignali; Addgene plasmid # 52114; http://n2t.net/addgene:52114; RRID:Addgene_52114) or the Retro-X™ Tet-One™ Inducible Expression System (Puro) (TakaRa Bio USA, Cat #634307). The *CSF3R* construct was cloned into a Gateway-converted pMSCV-IRES-GFP vector (a gift from Tannishtha Reya; Addgene plasmid #20672; http://n2t.net/addgene:20672; RRID:Addgene_20672). Plasmid sequences were confirmed via Sanger sequencing (Eurofins Genomics).

### Retrovirus generation and transduction

The human embryonic kidney 293T17 cell line (ATCC) was grown in DMEM (Gibco) medium with 10% fetal bovine serum (FBS, HyClone), Glutamax (Gibco), and penicillin/streptomycin (Gibco). Retrovirus was generated by transfecting *SETBP1* pMSCV-IRES-mCherry or *CSF3R* pMSCV-IRES-GFP or control vector plasmids and the EcoPac plasmid (provided by Dr. Rick Van Etten) into the 293T17 cells using FuGENE 6 (Promega). Viral supernatants were harvested 48 and 72 hours later. Isolated mouse bone marrow was cultured overnight in the presence of SCF, IL6, and IL3. Mouse bone marrow (1×10^6^ cells/well) was spinoculated with viral supernatant, HEPES buffer, and polybrene on two subsequent days. For the spinoculation, cells were spun at 2,500 rpm for 90 minutes at 30°C (brake turned off).

### Murine hematopoietic colony forming unit (CFU) assays

CFU assays were performed as described previously^1^. Retrovirus was generated as described above for *SETBP1, CSF3R, MYC* or control vector and used to transduce murine bone marrow cells isolated from 6- to 10-week-old female C57BL/6J mice, and cells positive for both constructs were sorted on a BD FACSAria III sorter into methylcellulose M3234 methylcellulose medium (StemCell Technologies). Cells were imaged using STEMvision (StemCell Technologies) and then manually counted using ImageJ (NIH). Images were blinded prior to counting by another investigator. For each replating, cells were isolated from methylcellulose, washed with PBS, and then plated in triplicate.

### Bone marrow transplantation 1

Bone marrow was harvested from 7-week-old female Balb/c mice and the mature hematopoietic cells were depleted using a Miltenyi Biotec Mouse Direct Lineage Cell Depletion Kit (130-090-858) with LS Columns (130-042-401) and 30µm Pre-Separation Filters (130-041-407). Retroviral-conditioned media was produced using 293T17 cells transduced with packaging plasmid and the appropriate transfer plasmid, as described above. Sorted double-positive cells were spun down and resuspended in PBS with non-transduced, fresh carrier bone marrow. Lethally irradiated Balb/c mice (2×4.5Gy) were transplanted with 25,000 transduced RFP+/GFP+ cells and 250,000 carrier bone marrow cells per mouse, via retro-orbital injection (N=7). Groups were randomly assigned. After transplantation, mice were maintained on antibiotic water for two weeks (Polymyxin B sulfate salt, Sigma-Aldrich P1004; NeoMycin trisulfate salt hydrate, Sigma-Aldrich N1876). Mice were monitored for onset of disease by investigators and animal care personnel.

### Bone marrow transplantation 2

Fluorouracil (100 mg/kg) was administered to 7-week-old female Balb/c mice (N=14) to deplete mature progenitor cells. Five days later, bone marrow was harvested from these mice. RBC were lysed using ACK lysis buffer and WBC were used for retroviral transduction. Retroviral-conditioned media was produced using 293T17 cells transduced with packaging plasmid and the appropriate transfer plasmid, as described above. Sorted double-positive cells were spun down and resuspended in PBS with non-transduced, fresh carrier bone marrow. Lethally irradiated Balb/c mice (2×4.5Gy) were transplanted with 2,000 transduced RFP+/GFP+ cells and 200,000 carrier bone marrow cells per mouse, via retro-orbital injection (N=6). Groups were randomly assigned. After transplantation, mice were maintained on antibiotic water for two weeks (Polymyxin B sulfate salt, Sigma-Aldrich P1004; NeoMycin trisulfate salt hydrate, Sigma-Aldrich N1876). Mice were monitored for onset of disease by investigators and animal care personnel.

### Histology

Cells shown in Figure 1C are representative images of cells contained within a single colony in the labelled CFU assay. Cells were harvested from single colonies using a Pasteur pipette and smeared onto a microscope slide. Dry slides were stained with May-Grünwald Stain (Sigma #63590) for two minutes and a 1:20 dilution of Giemsa stain for 12 minutes (Sigma-Aldrich #GS500). Representative cytospin preparations shown in 2F were prepared using Double Cytology Funnels (Fisherbrand #10-356) and stained with May-Grünwald and Giemsa as described above.

### Cell Line Generation

As previously described^12^, *CSF3R*^*T618I*^+*SETBP1*^*Wt*^ and *CSF3R*^*T618I*^+*SETBP1*^*D868N*^ murine cell lines were initially derived from mouse bone marrow cells that were harvested from CFU assays (Figure 1A-D). Subsequent cell lines, including the doxycycline-inducible cell line in Figure 3 (*CSF3R*^*T618I*^*+SETBP1*^*D868N*-dox^ cell line) were generated by sorting transduced murine bone marrow directly into IMDM with 20% FBS, without cytokine support. Cell lines were tested for Mycoplasma contamination monthly while in culture, before freezing and after thawing using a Lonza MycoAlert Mycoplasma Detection Kit (Catalog #: LT07-118).

### Flow Cytometry

For differentiation analyses, one million cells were resuspended in FACS buffer and stained with Cd11b PE-Cy7 (BD 552850, clone M1/70, lot 0058856) and GR-1 (Ly-6G and Ly-6C, BD 553128, clone RB6-8C5 (RUO), lot 9098937) for one hour. For cell cycle analyses, one million cells were fixed with ethanol, treated with RNase and stained with Propidium Iodide Staining Solution (BD 556463) for 30 minutes. Cells were analyzed using a BD FACSAria III, FlowJo (10.7.2) and FSC Express 7 Research software.

### RNA-seq

Doxycycline was withdrawn from the *CSF3R*^*T618I*^*+SETBP1*^*D868N*-dox^ cell line by washing the cells with PBS five times and then resuspending the cells in triplicate +/- 1 μM doxycycline. RNA was extracted from cells at 24h post-withdrawal using the RNeasy micro kit (Qiagen). cDNA libraries were constructed using the Takara SmartSeq for Ultra Low Input kit and sequenced using a HiSeq 2500 Sequencer (Illumina) 100 bp SR. Raw reads were trimmed with Trimmomatic^24^ and aligned with STAR^25^. Differential expression analysis was performed using DESeq2^26^. Raw p-values were adjusted for multiple comparisons using the Benjamini–Hochberg method. Enrichr^27,28^ and GSEA^29,30^ were used to assess gene set enrichment. Motif enrichment analysis was performed for DE genes using HOMER^13^.

### CUT&Tag

Doxycycline was withdrawn from the *CSF3R*^*T618I*^*+SETBP1*^*D868N*-dox^ cell line by washing the cells with PBS five times and then resuspending the cells in triplicate +/- 1 μM doxycycline. CUT&Tag methods were performed as previously described^31^. Histone CUT&Tag datasets were analyzed using an in-house CUT&Tag pipeline, script available upon request. Reads were aligned to the mm10 genome using Bowtie2^32^ and duplicates were marked but not removed with Sambamba^33^. Peaks were called using the CUT&Tag peak caller GoPeaks. Bedtools^34^ helped subset for peaks that appear in >= 2 replicates, which were used to build the feature count table and generate heatmaps. Differential open chromatin regions were identified using DESeq2^26^ with default settings, and then analyzed with HOMER motif analysis^13^. All differential peaks were intersected with MYC (ENCFF152JNC) and MYB (ENCFF911NHJ) ChIP-seq data from ENCODE^13^ using a murine erythro-leukemia cell line. The unique overlapping regions between differential CUT&Tag peaks and transcription factor binding sites were annotated using ChIPSeekeR^35^ and visualized with ggplot2^36^.

### Promoter Assay

Plasmids were transiently transfected into 293T17 cells using FuGENE 6 (Promega). For these experiments pGL2M4-luc reporter plasmid^37^ (containing four CACGTG binding sites, a canonical E-box) and *pRL Renilla Luciferase Control Reporter Vectors* (CMV, Promega E2231) were used. Luciferase activity was quantified using the Promega Dual-Luciferase Reporter Assay System (E1910) with a BioTek Synergy2 plate reader.

### Inhibitor Screening and Synergy Analysis

A mid-throughput chemical screen was performed as described previously^38^. An IC50 was calculated for each inhibitor based on the 7-point dose response curve (10-0.014 μM). This IC50 was expressed relative to the median IC50 for previously screened cell lines and human samples, thus allowing for generation of a measure of the fold efficacy^38^. Synergy analysis between ruxolitinib and LSD1 inhibitors as reported in **Figure 7** was performed using a drug dose matrix. The *CSF3R*^*T618I*^+*SETBP1*^*D868N*^ cell line was plated in an 8×8 matrix in triplicate with increasing concentrations of each inhibitor. Viability of cells assessed at 72 hours post-drug treatment using a tetrazolamine-based (MTS) assay, and synergy calculated with SynergyFinder^39^. Synergy in **Figure S6** was calculated by Bliss additivity analysis^40^. To determine if there was Bliss additivity, the *CSF3R*^*T618I*^+*SETBP1*^*D868N*^ cell line was plated in 96 well format and cells were treated for 72 hours with GSK2879552, ruxolitinib or the combination and viability of cells assessed using a tetrazolamine-based (MTS) assay. A value for Bliss additivity (predicted viability if the compounds had no synergy or antagonism) was calculated for each dose based on the single agent treatments, as previously described^40^. Synergy was assessed by assessing whether the predicted viability was greater or less than the viability of the combination treatment.

### qPCR

The cell line where *SETBP1*^D868N^ expression is regulated by doxycycline was treated with GSK2879552 (1000nM), GSK-LSD1 (100nM) or JQ1 (200nM) for 48 hours and qPCR was performed for *Myc*. In a second experiment, *CSF3R*^*T618I*^+*SETBP1*^*D868N*^ cell line with one of three LSD1 inhibitors at 100 nM (GSK2879552) or 30 nM (GSK-LSD1, ORY-1001) for 48 hours and qPCR was performed for *Myc, Myb, Meis1* and *Hoxa9*. RNA was extracted from cells using the RNeasy mini kit (Qiagen). cDNA was synthesized using the High-Capacity cDNA Reverse Transcription Kit with RNase Inhibitor (ThermoFisher). Taqman probes (ThermoFisher Scientific): *GusB* (Mm00446953_m1), *Myc* (Mm00487804_m1), *Myb* (Mm00501741_m1), *Meis1* (Mm00487664_m1) and *Hoxa9* (Mm00439364_m1). qPCR Reagents: TaqMan™ Universal PCR Master Mix (ThermoFisher). Instruments: T100 Thermal Cycler (BioRad), QuantStudio™ 7 Flex Real-Time PCR System (ThermoFisher).

### CNL patient sample processing and expansion

Live CD34+ progenitor cells were isolated from a CNL bone marrow sample with *CSF3R*^*T816I*^ and *SETBP1*^*G870S*^ mutations using Miltenyi Biotec Dead Cell Removal Kit (130-090-101) with LS columns (130-042-401) and EasySep™ Human CD34 Positive Selection Kit II (Catalog # 17856) with EasyEights™ EasySep™ Magnet (Catalog # 18103). The CD34+ cells were then cultured in a serum-free expansion media for 7 days (StemSpan™ SFEM II, Catalog #09605; StemSpan™ CD34+ Expansion Supplement, Catalog # 02691; UM729 Pyrimido-indole derivative that enhances HSC self-renewal in vitro, Catalog # 72332). The total number of CD34+ cells expanded from 65,400 to 642,000 cells in 7 days. 300,000 cells were then treated with either 100 nM ORY-1001 or DMSO for 24 hours. After treatment, single cell RNA-seq with barcoded antibody labeling (CITE-seq) was performed.

### CITE-Seq

Following treatment, cells were suspended in FACS staining buffer and incubated for 30 minutes with the following CITE-Seq TotalSeq A anti human primary antibodies: CD45RA (biolegend 304157), CD123 (biolegend 306037), CD10 (biolegend 312231), CD90 (biolegend 328135), CD34 (biolegend 343537), CD38 (biolegend 356635), CD366/Tim-3 (biolegend 345047), CD99 (biolegend 371317). A Chromium Single Cell 3 GEM, Library & Gel Bead Kit v3 (10X Genomics, PN-1000075) was utilized for the following steps. Cells were loaded onto the Chromium Controller (10X Genomics) according to the manufacturer’s instructions. After the post GEM-RT cleanup, ADT (Illumina TruSeq Small RNA Indices A: RPI1, RPI2) and HTO (Index plate T, Set A, PN-1000213/PN-2000240) additive primers were added to the RT reaction according to the manufacturer’s instructions. RNA and ADT/HTO libraries were separated using Quantabio sparQ PureMag Beads and libraries were prepared separately according to the manufacturer’s instructions. Libraries were sequenced on a NexSeq system (Illumina) with a High Output Kit, 150 Cycles. CITE-Seq reads and feature barcodes were aligned to the pre-built single cell GRCh38 2020-A reference index from 10X Genomics using CellRanger v5.0.1. Secondary analysis was facilitated with a custom pipeline (https://github.com/maxsonBraunLab/cite_seq) using R v4.0.3 and Seurat v4.0.0^41^. Briefly, each sample was imported into Seurat, then filtered for UMI counts between 2K – 20K and mitochondrial content < 15%. Cell cycle regression was applied to each sample before integration with DMSO as the baseline condition using 50 PCs integration anchor and k anchor of 5. The individual samples and the integrated object were clustered with UMAP with 50 dimensions before manual annotation of cell types and visualization of MYC expression per cluster per condition.

### Data presentation

All graphs were made using either ggPlot2, GSEA or GraphPad Prism; figures were assembled in Adobe Illustrator. Data is presented as mean ± standard error of the mean.

### Data sharing and key resources

Contact Julia Maxson at for information regarding renewable materials, datasets, and protocols. Sequencing data will be available on GEO following peer review.

## Acknowledgements

Research reported in this publication was supported by: an American Society of Hematology Research Restart Award, Collin’s Medical Trust Award, Medical Research Foundation Early Clinical Investigator Award, and NCI F32CA239422 to SAC; an American Society of Hematology Research Restart Award, an American Society of Hematology Scholar Award and 1 K08 CA245224 from NCI awarded to TPB; a Knight Pilot Award, American Society of Hematology Junior Faculty Scholar Award, Gilead Research Scholars Program in Hematology/Oncology, and 1R01HL157147-01 to JEM. We thank the following OHSU core facilities for their assistance: Advanced Light Microscopy, Flow Cytometry Shared Resource, ExaCloud Cluster Computational Resource, and the Advanced Computing Center.

## Author Contributions

*Concept and design:* Carratt, Druker, Braun, Maxson

*In vitro experiments:* Carratt, Smith, Baris, Braun

*In vivo experiments:* Carratt, Schonrock, Maloney

*Computational resources and analysis:* Kong, Smith, Braun

*Analysis and interpretation of data:* Carratt, Kong, Smith, Druker, Braun, Maxson

*Writing, review and revision of the manuscript:* All authors

---

**Figure S1.**
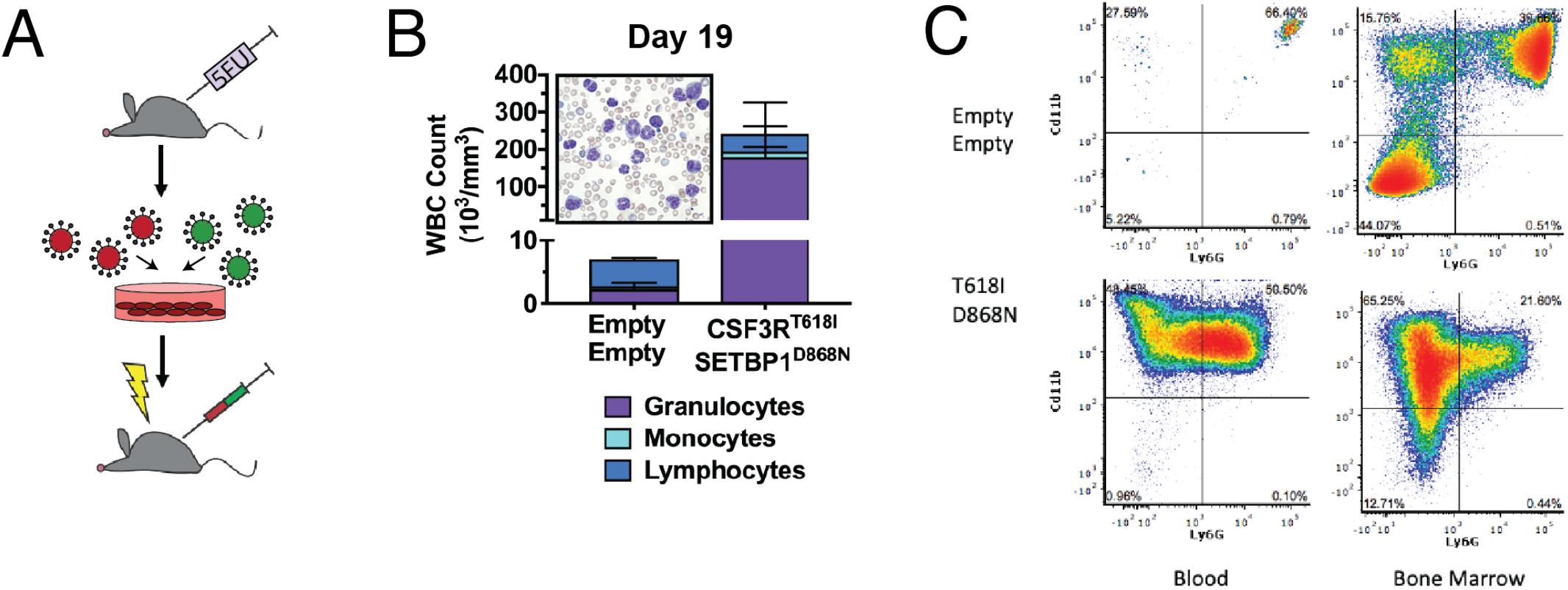
Expansion of the granulocyte lineage following transplantation of transduced bone marrow. **(A)** Schematic for transplant model. Balb/c donors were treated with Fluorouracil (5FU) via retro orbital injection (100 mg/kg) six days prior to bone marrow harvest. Marrow from donor mice was transduced with *CSF3R*^*T618I*^ and *SETBP1*^*D868N*^, or with empty vector controls. Cells positive for both *CSF3R* and *SETBP1* were sorted and 2,000 cells per condition—along with 200,000 carrier marrow cells—were transplanted into lethally irradiated mice. Controls were transplanted with 2,000 cells double positive for the empty vectors. **(B)** At Day 19, mice transplanted with *CSF3R*^*T618I*^+*SETBP1*^*D868N*^ cells (N=3) had high white blood cell counts. A representative peripheral blood smear from these mice shows a prevalence of mature myeloid cells (*insert*). **(C)** Flow cytometry for Cd11b and Ly6G in the *CSF3R*^*T618I*^+*SETBP1*^*D868N*^ mice shows an abundance of Cd11b+ cells in the peripheral blood and an increase in the Cd11b+/Ly6G-cells in the bone marrow compartment.

**Figure S2.**
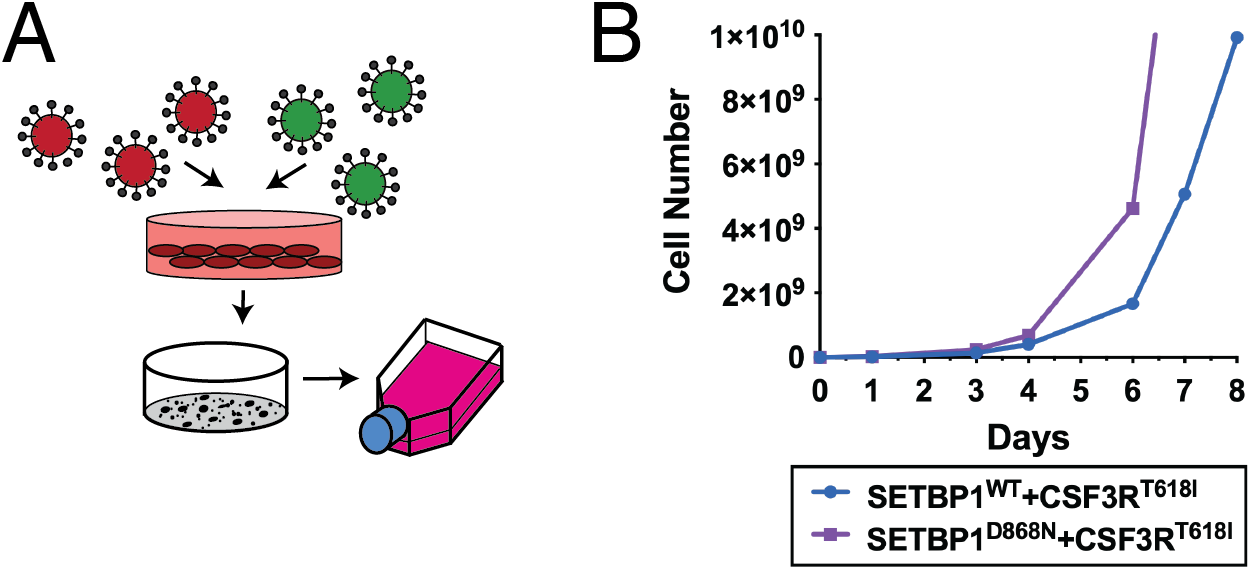
Mouse bone marrow cells immortalized with *SETBP1* and *CSF3R*. **(A)** Mouse bone marrow cells were harvested from CFU assays (**Figure 1**). Cells were washed and maintained in liquid culture (IMDM with 20% FBS) without any additional cytokine support. Within one week, both *CSF3R*^*T618I*^+*SETBP1*^*Wt*^*and CSF3R*^*T618I*^+*SETBP1*^*D868N*^ transduced cells began to grow exponentially. Cells were maintained in liquid culture in the absence of cytokines and continued to proliferate, thus generating a cell line. **(B)** Cell line growth over 8 days.

**Figure S3.**
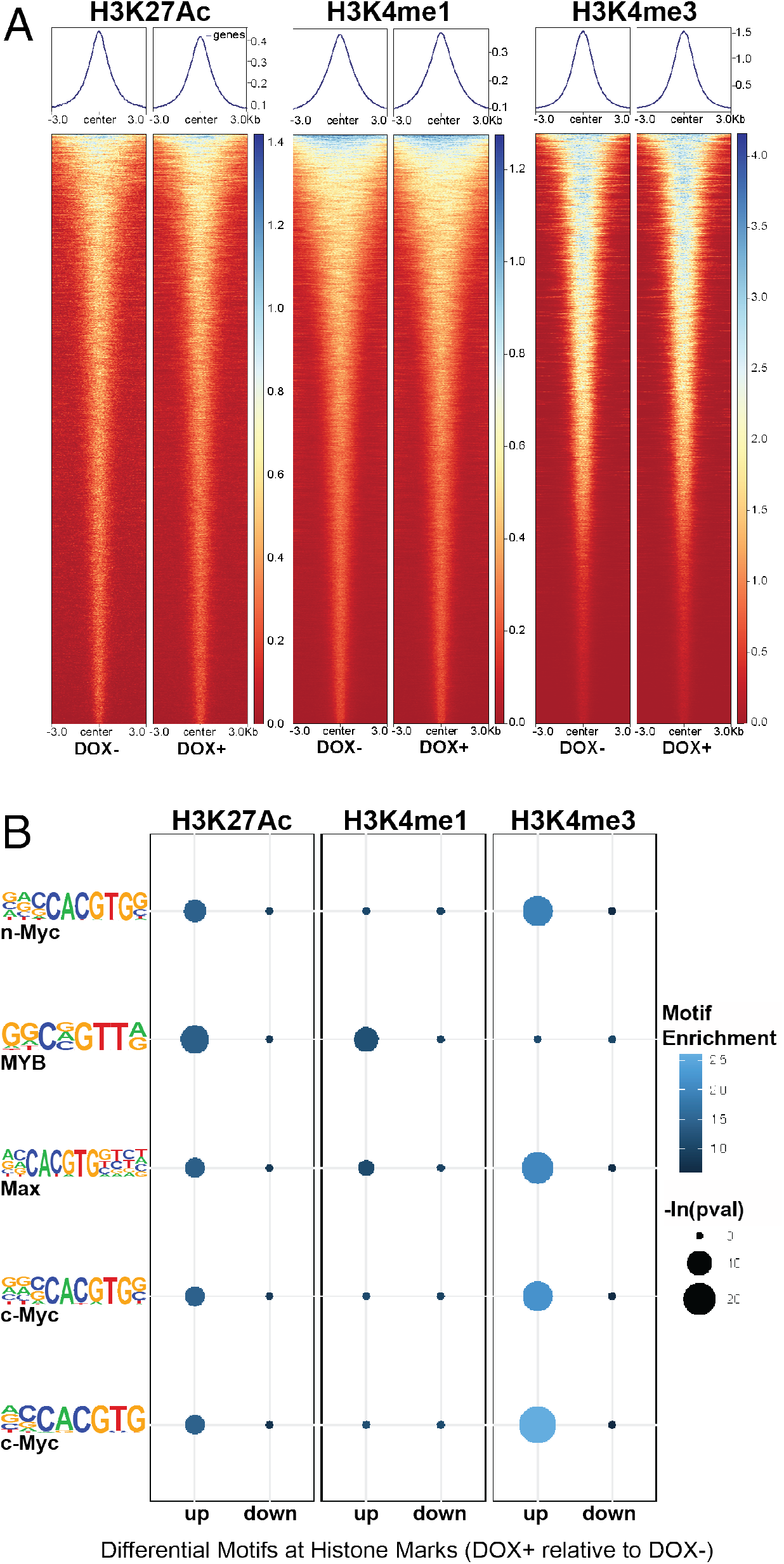
CUT&Tag for histone marks. **(A)** Average shape of the peaks at consensus intervals. There is not a global decrease in deposition of epigenetic marks. **(B)** MYC motifs identified for each histone mark at differential peaks between DOX+ and DOX-conditions. Motif enrichment shown for DOX+ relative to DOX-.

**Figure S4.**
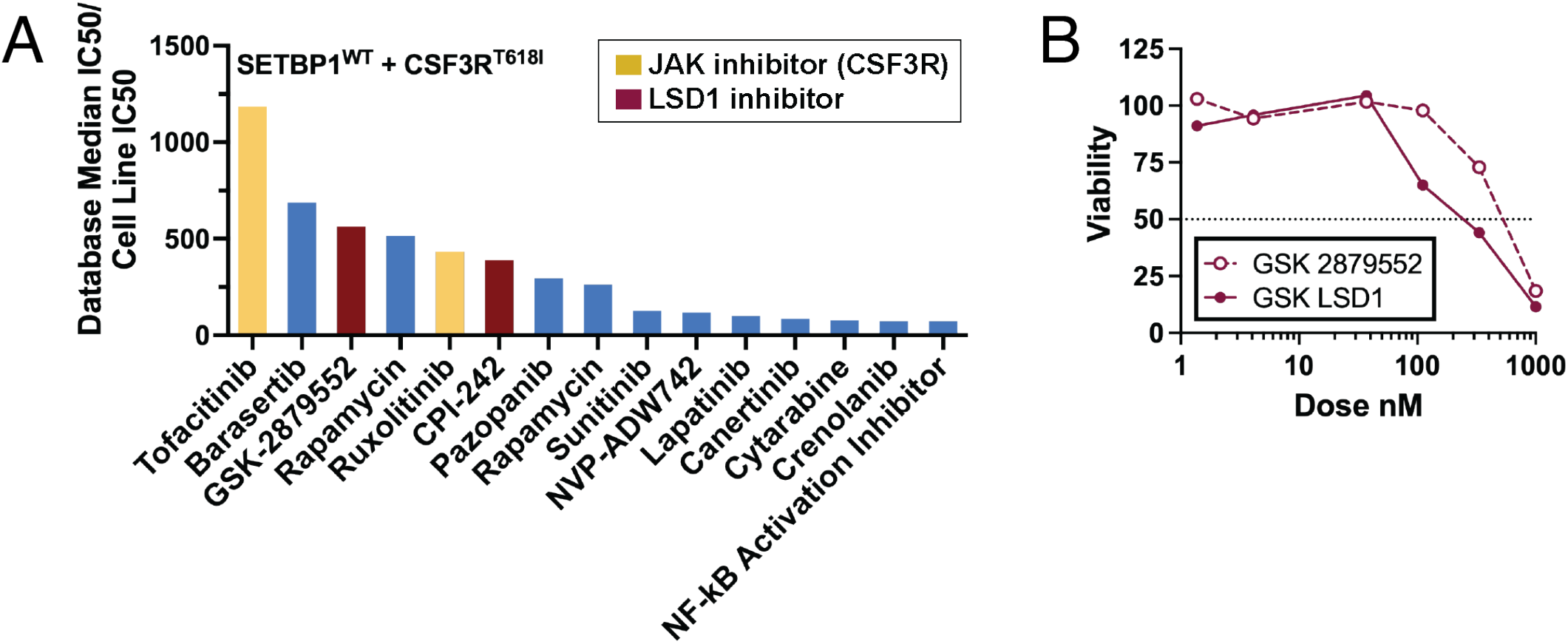
Inhibitor sensitivity data. **(A)** Using the *CSF3R*^*T618I*^+*SETBP1*^*Wt*^ cell line, which was generated by retrovirally transducing *CSF3R*^T168I^ and *SETBP1*^*Wt*^ into mouse bone marrow, we performed a screen for dependencies as described for the SETBP1^D868N^ cell line. When a chemical screen with 175 inhibitors with known sensitivity in patient samples (BeatAML cohort^14^) was performed on this wildtype cell line, these cells were also sensitive to both JAK and LSD1 inhibitors. **(B)** Dose response curve for two LSD1 inhibitors (1-1000 nM) in the *CSF3R*^*T618I*^+*SETBP1*^*D868N*^ cell line. The IC50s are approximately 590 nM (GSK2879552) and 250 nM (GSK-LSD1).

**Figure S5.**
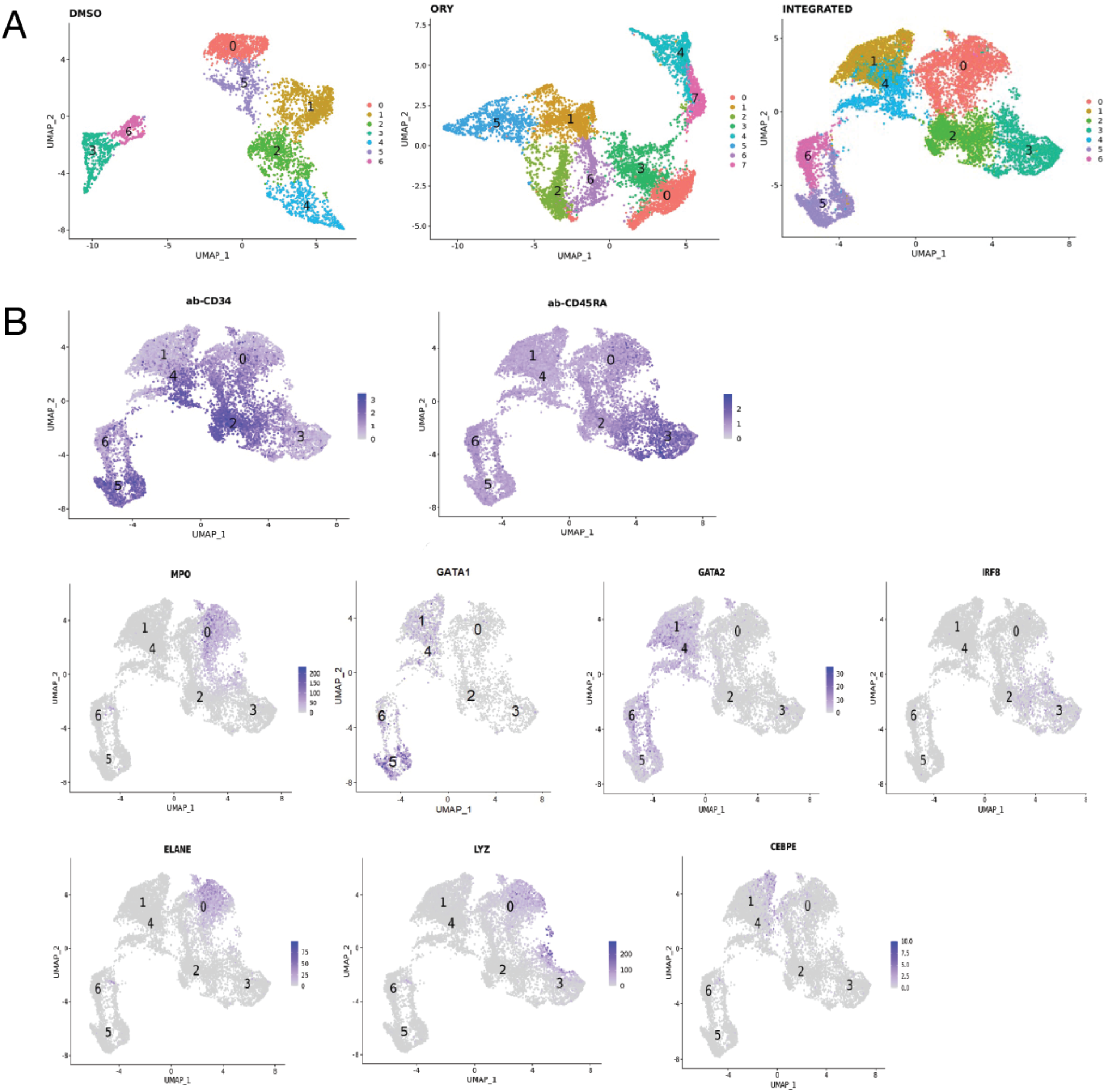
Identification of population hierarchy in a patient sample treated with DMSO or ORY-1001. Cryopreserved bone marrow from a chronic neutrophilic leukemia patient was thawed and live CD34+ cells were cultured for 7 days in a serum-free CD34 expansion medium (STEMCELL StemSpan SFEM II with CD34+ Expansion Supplement and UM729). Between days 7-8, cells were treated for 24 hours with either DMSO or ORY-1001 (N=1) and Cellular Indexing of Transcriptomes and Epitopes by Sequencing (CITE-seq) was performed. **(A)** Uniform Manifold Approximation and Projection (UMAP) clustering of samples individually and clustering following sample integration. **(B)** Marker genes and surface antigens used for population identification.

**Figure S6.**
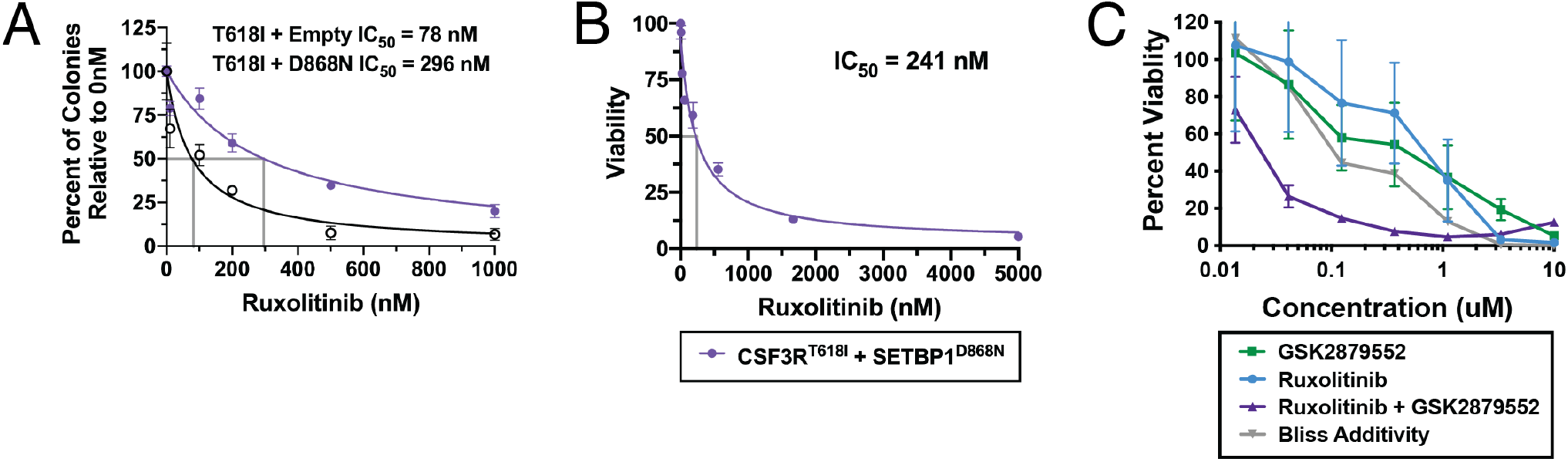
Ruxolitinib sensitivity in *SETBP1*^*D868N*^ mutant cells. **(A)** CFU assay showing sensitivity to ruxolitinib for cells transduced with *CSF3R*^*T618I*^+empty vector or *CSF3R*^*T618I*^+*SETBP1*^*D868N*^. Primary mouse bone marrow was harvested from C57BL/6 mice and retrovirally transduced with either *CSF3R*^*T6181*^+empty vector or *CSF3R*^*T6181*^+*SETBP1*^*D868N*^. Double positive cells were sorted into methylcellulose and a cytokine-free colony unit forming (CFU) assay was performed with 4 doses of ruxolitinib (100nM, 500nM, 200nM, 100nM). Because of the different baseline between the number of colonies formed, 30,000 cells were plated for the *CSF3R*^*T618I*^+empty vector conditions and 5,000 for the *CSF3R*^*T618I*^+*SETBP1*^*D868N*^ conditions, resulting in a baseline of approximately 40 and 115 CFUs at 0nM, respectively. Colonies formed are reported as a percentage of control and IC50 was estimated assuming a standard slope equal to a Hill slope of -1.0. **(B)** The IC50 of the *CSF3R*^*T618I*^+*SETBP1*^*D868N*^ cell line is 241 nM. The *CSF3R*^*T618I*^+*SETBP1*^*D868N*^ immortalized mouse bone marrow cell line was treated with ruxolitinib using a 1:3 serial dose response curve (7-5000 nM). Viability was measured at 72 hours with a tetrazolamine-based (MTS) assay. **(C)** Bliss adaptivity analysis of GSK2879552 and ruxolitinib in the *CSF3R*^*T618I*^+*SETBP1*^*D868N*^ cell line. To evaluate if there was synergy between the LSD1 and JAK inhibition in the *CSF3R*^*T618I*^+*SETBP1*^*D868N*^ cell line, cells were treated for 72 hours with GSK2879552, ruxolitinib or the combination. Bliss additivity (grey) indicates the predicted viability if the compounds had no synergy or antagonism. The combination (purple) viability values fell far below this line, indicating synergy between GSK2879552 and ruxolitinib.

**Table S1.** Annotations for top inhibitors of the *CSF3R*^*T618I*^*+SETBP1*^*D868N*^ cell line.

| **Drug** | **Target** | **Fold change over median IC50** |
| --- | --- | --- |
| *CPI-242* | LSD1 | 835 |
| *GSK-2879552* | LSD1 | 421 |
| *Barasertib* | Aurora B | 403 |
| *Rapamycin* | mTOR | 226 |
| *Crenolanib* | pan-FLT3 | 124 |
| *AT-101* | pan-Bcl-2 | 121 |
| *Ruxolitinib* | JAK | 75 |
| *Selumetinib* | MEK/ERK | 62 |
| *Cytarabine* | antimetabolite | 57 |
| *NF-κB Activation Inhibitor* | NF-κB | 52 |
| *JAK Inhibitor I* | JAK | 41 |
| *Lapatinib* | EGFR, HER-2 | 41 |
| *Artemisinin* | anti-malarial | 34 |
| *Tozasertib (VX-680)* | pan-Aurora | 32 |
| *Pelitinib (EKB-569)* | EGFR | 22 |

**Table S2.** Annotations for top inhibitors of the *CSF3R*^*T618I*^+*SETBP1*^*Wt*^ cell line.

| **Drug** | **Target** | **Fold change over median IC50** |
| --- | --- | --- |
| *Tofacitinib* | JAK3 | 1185 |
| *Barasertib* | Aurora B | 688 |
| *GSK-2879552* | LSD1 | 564 |
| *Rapamycin* | mTOR | 515 |
| *Ruxolitinib* | JAK | 433 |
| *CPI-242* | LSD1 | 390 |
| *Pazopanib* | VEGFR-1, VEGFR-2, VEGFR-3, PDGFR-α/β, and c-kit | 295 |
| *Rapamycin* | mTOR | 262 |
| *Sunitinib* | multi-target RTK | 127 |
| *NVP-ADW742* | IGF-1R | 118 |
| *Lapatinib* | EGFR, HER-2 | 100 |
| *Canertinib* | EGFR, HER-2 and ErbB-4 | 85 |
| *Cytarabine* | antimetabolite | 77 |
| *Crenolanib* | pan-FLT3 | 72 |
| *NF-κB Activation Inhibitor* | NF-κB | 72 |

